# A feed-forward UHRF1 read-write mechanism supports H3 multi-mono-ubiquitination and DNA methylation maintenance at CpG-sparse regions

**DOI:** 10.64898/2026.09.25.754493

**Authors:** Joel A. Hrit, Rochelle L. Tiedemann, Vincent J. Sartori, Ashley K. Wiseman, Bradley M. Dickson, Evan J. Worden, Scott B. Rothbart

## Abstract

The epigenetic inheritance of mammalian DNA methylation requires DNMT1 and its E3 ligase cofactor UHRF1. At newly replicated chromatin, UHRF1 recognition of hemi-methylated DNA and histone H3 N-terminal tails directs catalysis of H3K14, H3K18, and/or H3K23 mono-ubiquitination to recruit DNMT1. While it is appreciated that UHRF1 can deposit multiple mono-ubiquitin marks on a single H3 tail and that DNMT1 recognizes this state through tandem ubiquitin interacting motifs, the mechanism that promotes successive ubiquitination and the biological function of multi-mono-ubiquitination are unknown. Here, we show that UHRF1 directly binds its mono-ubiquitinated H3 products through a previously uncharacterized LGDDSL loop in Tudor 2 of its tandem Tudor domain (TTD) to promote further ubiquitin deposition. Disruption of this ubiquitin reading activity impairs H3 multi-mono-ubiquitination and accelerates DNA methylation loss within late-replicating, CpG-sparse genomic regions that are characteristic of partially methylated domains (PMDs) in cancer and aging cells. These methylation defects overlap those observed by disruption of UHRF1 ubiquitin ligase activity, providing convergent evidence that both writing and reading of H3 ubiquitination support CpG-sparse DNA methylation maintenance. Together, these findings establish a feed-forward ubiquitin read-write mechanism that generates multi-mono-ubiquitinated H3 and safeguards DNMT1-dependent DNA methylation maintenance at vulnerable genomic regions of the mammalian methylome.

## INTRODUCTION

DNA methylation is an essential epigenetic regulator of mammalian development, contributing to the establishment and maintenance of transcriptional programs that define cellular identity^1–3^. DNA methylation patterns are established in early embryonic development by *de novo* methyltransferases and inherited through somatic cell divisions by the maintenance DNA methyltransferase DNMT1^4^. Disruption of these patterns is an epigenetic hallmark of nearly all human cancers and a major contributor to oncogenesis and tumor progression^5^. In normal cells, dense clusters of unmethylated CpG dinucleotides (CpG islands) are enriched at promoters and regulatory elements, whereas CpGs sparsely distributed throughout gene bodies and the intergenic genome are heavily methylated^6,7^. This organization is disrupted in cancer cells, where CpG island promoter hypermethylation contributes to tumor suppressor gene silencing and suppression of tumor immunogenicity, while widespread hypomethylation of intergenic DNA promotes genome instability and retrotransposition^5,8–10^. Notably, DNA hypomethylation is particularly pronounced within late-replicating, CpG-sparse regions of the genome, where progressive methylation loss gives rise to partially methylated domains (PMDs) in cancer and aging cells^11,12^. While accurate propagation of DNA methylation patterns is essential for preserving epigenome integrity, the mechanisms that support their inheritance through mitotic cell divisions, and the extent to which disruption of these processes contributes to the progressive methylation loss associated with cancer and aging, remain incompletely understood.

DNA methylation inheritance is coupled to the process of semi-conservative DNA replication^13–15^. In the wake of replicating DNA polymerase, hemi-methylated DNA (heDNA) intermediates form from the annealing of parental methylated DNA to nascent unmethylated DNA. DNMT1 catalyzes methylation of these heDNA intermediates to reform symmetrically methylated DNA^15,16^. The recruitment of DNMT1 to its substrates is facilitated by the E3 ubiquitin ligase UHRF1^13,14,17–19^. This stepwise mechanism involves UHRF1 chromatin targeting through coordinated activities of its tandem Tudor domain (TTD) that binds lysine 9 di- or tri-methylation on histone H3 (H3K9me2/me3)^18–21^, its plant homeodomain (PHD) finger that binds the unmodified N-terminus of H3^18,19,21,22^, and its SET- and RING-associated (SRA) domain that binds selectively to heDNA^23–25^. These multivalent substrate targeting events direct its UBL- and RING domain-dependent enzymatic activity toward ubiquitination of histone H3 on lysines 14, 18, and/or 23^17,26–28^. DNMT1 has tandem ubiquitin interacting motifs (UIMs) embedded in its replication foci targeting sequence (RFTS) that recognize the H3 ubiquitination sites that UHRF1 installs^29–33^. This elegant ‘read-write-read-write’ mechanism couples histone H3 ubiquitination to the epigenetic inheritance of DNA methylation UHRF1 can generate mono- and multi-mono-ubiquitination on a single H3 tail both *in vitro* and in cells^17,28,34^. Ubiquitination of H3K14, K18, or K23 (alone or in combination) creates binding sites for the DNMT1 RFTS domain^32^, and structural analyses and quantitative binding studies demonstrate that the tandem UIMs (UIM1 and UIM2) of the RFTS domain engage in multivalent interactions with multi-mono-ubiquitinated H3 peptides^32^. Recognition of H3 ubiquitination by the tandem UIMs is indispensable for DNA methylation maintenance^31,35^.

However, whether multi-mono-ubiquitination of H3 is strictly required for maintenance DNA methylation in cells remains unresolved. Notably, our prior work showed that disruption of either DNMT1 UIM1 or UIM2 alone had a more modest effect on DNA methylation maintenance than their combined disruption^35^, suggesting that the requirement for multi-mono-ubiquitin recognition may be context dependent. Moreover, while the PHD and SRA domains are necessary for UHRF1 chromatin recruitment and E3 ligase activity, the mechanisms that promote successive ubiquitin deposition on an individual H3 tail remain unknown.

In this study, we identify a previously unrecognized interaction between UHRF1 and its mono-ubiquitinated H3 products that promotes further ubiquitin deposition and the generation of multi-mono-ubiquitinated H3 both *in vitro* and in cells. We map this ubiquitin reading activity to a previously uncharacterized loop within Tudor 2 of the UHRF1 TTD and show that, analogous to H3K9me2/me3 recognition by Tudor 1, H3ub recognition by Tudor 2 is anchored through PHD finger engagement of the H3 N-terminus. Disruption of UHRF1 ubiquitin reading activity in colon cancer cells reduces H3 multi-mono-ubiquitination and accelerates DNA methylation loss during serial passaging, with preferential effects at late-replicating, CpG-sparse genomic regions that overlap those affected by disruption of UHRF1 ubiquitin ligase activity and are susceptible to the formation of PMDs in cancer and aging cells. Collectively, these findings reveal how UHRF1 product recognition promotes successive ubiquitin deposition and establish a functional requirement for multi-mono-ubiquitination in preserving DNA methylation within these vulnerable genomic regions.

## MATERIALS AND METHODS

### Reagents

Commercial kits used for this study included: QIAprep Spin Miniprep kit (Catalog # 27 104) and QIAquick PCR purification kit (Catalog # 28104, Qiagen, Germantown, MD, USA), High Sensitivity Qubit Fluorometric Quantification (Catalog # Q32854, Invitrogen, Carlsbad, CA, USA), Alpha Screen Histidine detection kit (Catalog # 6760619M, Revvity, Waltham, MA, USA), and BCA protein assay kit (Catalog # 23227, Thermo Fisher, Waltham, MA, USA).

Antibodies used for western blotting included: DNMT1 (Catalog # ab134148, Abcam, MA, USA), UHRF1 (Catalog # D6G8E, Cell Signaling Technology, Danvers, MA, USA), H3K18ub (Catalog # E4D7R, Cell Signaling Technology, Danvers, MA, USA), H3K9me3 (Catalog # 39161, Active Motif, Carlsbad, CA, USA), histone H3 (Catalog # 13-0001, Epicypher, Durham, NC, USA), and rabbit IgG HRP-conjugated secondary antibody (Catalog # GENA934, Sigma-Aldrich, Burlington, MA, USA).

Non-standard chemicals and reagents used in the study included: Recombinant histones H2A, H2B, H3, and H4 (Catalog # XH2A, XH2B, XH3, and XH4 respectively, The Histone Source, Fort Collins, CO, USA), His60 Ni Superflow Resin (Catalog # 635662, Takara Bio USA, San Jose, CA, USA), Glutathione Agarose Resin (Catalog # G-250-100, GoldBio, St. Louis, MO, USA), Benzonase Nuclease (Catalog # E1014, Millipore Sigma, Burlington, MA, USA), Lysozyme (Catalog # 89833, Thermo Fisher Scientific, Waltham, MA, USA), 1,3-dichloroacetone (Catalog # A14149.14, Thermo Fisher Scientific, Waltham, MA, USA), RNAse A/T1 mix (Catalog # EN0551, Thermo Fisher Scientific, Waltham, MA, USA), Proteinase K (Catalog # 25530015, Thermo Fisher Scientific, Waltham, MA, USA), Streptavidin Sepharose beads (Catalog # 17-5113-01, Cytiva, Marlborough, MA, USA), Glutathione Magnetic Agarose beads (Catalog # 78602, Thermo Fisher Scientific, Waltham, MA, USA), doxycycline (dox) (hyclate) (Catalog # 14 422, Cayman Chemical, Ann Arbor, MI, USA), 16% methanol-free formaldehyde solution (Catalog # 28 906, Thermo Fisher Scientific, Waltham, MA, USA), X-tremeGENE HP DNA Transfection Reagent (Catalog # XTGHP-RO, Roche, Indianapolis, IN, USA), Protein Assay Dye Reagent Concentrate (Catalog # 5 000 006, Bio-Rad, Hercules, CA, USA), AlphaLISA GSH acceptor beads (Catalog # AL109M, Revvity, Waltham, MA, USA), Alpha Streptavidin donor beads (Catalog # 6760002, Revvity, Waltham, MA, USA), AlphaPlate 384-well, light grey (Catalog # 6057350, Revvity, Waltham, MA, USA) and TAMRA-ubiquitin (Life Sensors, Malvern, PA, USA).

FPLC chromatography columns used for DNA and protein purifications in this study included: 1mL HisTrap HP (Catalog # GE17-5319-01), HiLoad 16/600 superdex 200 (Catalog # GE28-9893-35), 6mL RESOURCE Q (Catalog # GE17-1179-01), 5mL HiTrap SP HP (Catalog # GE17-1152-01), 5mL HiTrap Q HP (Catalog # GE17-1154-01), and HiPrep 26/60 Sephacryl S-200 HR (Catalog # 17-1195-01, Cytiva, Marlborough, MA, USA). HPLC chromatography columns included: PROTO 300 C4 5 micron, 250×10mm (Catalog # RS-0546-W045, Higgins Analytical, Mountain View, CA, USA).

Specialized commercial chips and instruments used in the study included: iScan system and Infinium MethylationEPIC BeadChIP v2.0 (Illumina, San Diego, CA, USA), and a Covaris E220 evolution (Covaris, Woburn, MA, USA).

### Biological Resources

Cell lines used in the study included HCT116 (CCL-247) and HEK 293T Phoenix AMPHO (CRL-3213) (ATCC, Manassas, VA, USA). Stable competent *Escherichia coli* (High Efficiency) (Catalog # C3040I, NEB, Ipswich, MA, USA) and XL10-Gold ultracompetent *Escherichia coli* (Catalog # 200315, Agilent, Santa Clara, CA, USA) were used for cloning. BL21 (DE3) Competent *E. coli* (Catalog # C2527I, NEB, Ipswich, MA, USA) and BL21 Rosetta2(DE3) Competent *E. coli* (Catalog # 71397, Novagen, Madison, WI, USA) were used for recombinant protein expression. Plasmids used in the study included: pMXs-IRES-blasticidin retroviral vector (Catalog # 72 876), pQE-80L for production of recombinant His-MBP-tagged proteins (Catalog # 32 943, Qiagen, Germantown, MD, USA), pGEX4T1 for production of recombinant GST-tagged proteins (Catalog # 28-9545-49, Cytiva, Marlborough, MA, USA), and pET3a for production of recombinant histone H3K18C and recombinant His-TEV-ubiquitin G76C (Catalog # 69418, Novagen, Madison, WI, USA).

### Recombinant protein expression and purification

His-MBP-tagged UHRF1 full length, GST-tagged UHRF1 TTD-PHD, and His-MBP-tagged DNMT1 RFTS were transformed into BL21(DE3) E. coli and grown at 37°C in LB + ampicillin. When OD reached 0.6-0.8, cultures were cooled to 16C and IPTG was added to 0.5mM (UHRF1 FL) or 0.2mM (TTD-PHD and RFTS). Cultures were grown at 16°C overnight to induce protein expression. Cells were harvested by centrifugation at 4,000rpm at 4°C for 20 minutes.

Cell pellets were resuspended in lysis buffer, flash frozen in liquid nitrogen, and stored at -80°C (UHRF1 FL: 50mM HEPES pH 7.5, 500mM NaCl, 20mM imidazole, 1mM TCEP, 1mM PMSF; UHRF1 TTD-PHD: 1x PBS, 1mM DTT, 1mM PMSF; DNMT1 RFTS: 50mM HEPES pH 7.5, 250mM NaCl, 20mM imidazole, 30μM ZnOAc, 0.5mM DTT, 1mM PMSF).

For purification of His-MBP-UHRF1 FL, lysis was performed by addition of lysozyme and incubation at 4°C with rotation for 30 minutes, followed by 3-4 rounds of sonication (10% amplitude, 1s on/1s off for 30s on, 1 min rest). Lysate was cleared by centrifugation at 38,000g at 4°C for 30 minutes. Soluble lysate was applied to a 1mL HisTrap column (Cytiva). The column was washed with 10-20 column volumes of wash buffer (50mM HEPES pH 7.5, 1M NaCl, 20mM imidazole, 1mM TCEP). Bound proteins were eluted with 10 column volumes of elution buffer (25mM HEPES pH 7.5, 100mM NaCl, 250mM imidazole, 1mM TCEP). Positive fractions were pooled and concentrated and UHRF1 was further purified by size exclusion chromatography using a Superdex 200 16/60 column in 25mM HEPES pH 7.5, 100mM NaCl, 1mM DTT. Positive fractions were pooled and concentrated by centrifugation using Amicon spin concentrators (50kDa MWCO). Glycerol was added to 10% final and protein was flash frozen in liquid nitrogen in small aliquots and stored at -80°C.

For purification of GST-TTD-PHD, cells were lysed and lysate was cleared as described above. Soluble lysate was added to glutathione Sepharose resin and incubated at 4°C with rotation for at least 1 hour. Resin was pelleted by centrifugation and washed 3-4 times with 10 bed volumes of lysis buffer. Bound protein was eluted in elution buffer (1x PBS, 1mM DTT, 10mM glutathione). Eluted protein was concentrated and exchanged into storage buffer (25mM HEPES pH 7.5, 100mM NaCl, 1mM DTT) by centrifugation using Amicon spin concentrators (30kDa MWCO). Glycerol was added to 10% final and protein was flash frozen in liquid nitrogen in small aliquots and stored at -80°C.

For purification of His-MBP-RFTS, cells were lysed and lysate was cleared as described above. Soluble lysate was added to His60 Ni superflow resin (Takara) and incubated at 4°C with rotation for at least 1 hour. Resin was pelleted by centrifugation and washed 3-4 times with 10 bed volumes of lysis buffer. Bound protein was eluted in elution buffer (25mM HEPES pH 7.5, 100mM NaCl, 250mM imidazole, 1mM TCEP). Eluted protein was concentrated by centrifugation using Amicon spin concentrators (30kDa MWCO). Glycerol was added to 10% final and protein was flash frozen in liquid nitrogen in small aliquots and stored at -80°C.

His-tagged ubiquitin G76C was transformed into BL21 Rosetta2(DE3) E. coli and grown at 37°C in LB + ampicillin + chloramphenicol. Cultures were grown and induced as above, except that IPTG was added to 1mM. Harvested cells were resuspended in lysis buffer (50mM Tris pH 7.8, 500mM NaCl, 250mM imidazole, 1mM PMSF, 5mM BME), flash frozen in liquid nitrogen and stored at -80°C. Cells were lysed as described above followed by addition of 10U DNase + 5mM MgCl_2_ and incubation at room temperature for 30 minutes. Lysate was cleared as described above, added to His60 Ni superflow resin (Takara), and incubated at 4°C for at least 1 hour. A gravity column was poured and washed with at least 10 bed volumes of lysis buffer.

Bound protein was eluted in elution buffer (50mM Tris pH 7.8, 500mM NaCl, 250mM imidazole, 1mM PMSF, 5mM BME) and dialyzed overnight into 4L of SP buffer A (50mM NH_4_OAc pH 4.5, 1mM DTT) at 4°C. Protein was further purified by ion exchange chromatography using 2×5mL HiTrap SP HP columns in series and a gradient from 0-1M NaCl. Positive fractions were pooled and dialyzed overnight into 4L of storage buffer (10mM Tris pH 7.8, 0.5mM TCEP) at 4°C. Protein was concentrated using spin concentrators (Thermo Scientific, 3kDa MWCO), flash frozen in small aliquots, and stored at -80°C.

Histone H3K18C/C110A was expressed and purified according to standard methods described previously^36^

### Preparation of E. coli soluble lysates for alpha screen

GST-TTD-PHD constructs were expressed in BL21(DE3) E. coli as described above. Cells were resuspended in lysis buffer (50mM HEPES pH 7.5, 50mM NaCl, 2mM MgCl_2_, 10% glycerol, 1mM PMSF, 1mM DTT, 2.5U/uL benzonase, lysozyme) and incubated at room temperature for 15 minutes to lyse. Lysate was cleared by centrifugation at 15,000rpm for 5 minutes. The concentration of overexpressed GST-TTD-PHD was estimated by running lysates and BSA standards on an SDS-PAGE gel and staining with Coomassie.

### Preparation of H3K18C-ub

H3K18C with a dichloroacetone linkage to ubiquitin G76C was prepared according to a protocol previously described for production of H2BK120C-ub^37^ with minor modifications. Briefly, H3K18C/C110A (100μM) and His-TEV-ubiquitin G76C (100μM) were crosslinked by addition of dichloroacetone (DCA) (100μM) and incubation on ice for 1 hour. Unreacted H3 and H3-H3 dimers were removed by Ni-NTA chromatography. The tag was cleaved off of ubiquitin with His-TEV protease, and uncleaved ubiquitin and His-TEV protease were removed by Ni-NTA chromatography. Finally, H3K18C-ub was purified from unreacted ub and ub-ub dimers by HPLC using a PROTO 300 C4 5 micron, 250×10mm column using a gradient of 0-90% acetonitrile in 0.1% TFA. Fractions containing pure H3K18C-ub were combined, lyophilized, and stored at -20°C.

### Preparation of ubiquitinated peptides

H3(1-20) peptides with or without trimethylation at K9, with K14C or K18C substitutions, were purchased from GenScript. Peptide (200μM) and His-TEV-ubiquitin G76C (100μM) were mixed in crosslinking buffer (50mM Borate pH 8.1, 1mM TCEP) and incubated to 50°C for 1 hour to reduce cysteines. Dichloroacetone (DCA) was dissolved in DMF and added to the reactions (100μM final concentration). Crosslinking was performed on ice for 1 hour, after which reactions were quenched by adding BME to 50mM final. Reactions were dialyzed overnight at 4°C into 4L of crosslinking buffer to remove unreacted peptide. Ubiquitin, ubiquitin dimers, and peptide dimers were removed by ion exchange chromatography using a 5mL HiTrap SP HP column (Cytiva) with a gradient elution from 0-1M NaCl in crosslinking buffer. Fractions containing ubiquitinated peptide were pooled, and the His6 tag was removed from ubiquitin by addition of His6-TEV protease and dialysis into 4L of TEV cleavage buffer (50mM Tris pH 8, 0.5mM EDTA, 1mM TCEP) overnight at 4°C. To remove His6-TEV and any remaining His6-tagged ubiquitin, samples were bound to Ni NTA resin (TakaraBio), and flowthru and washes containing pure ubiquitinated peptide were collected and pooled. Samples were concentrated by centrifugation using Amicon spin concentrators (3kDa MWCO), dialyzed into water, and flash frozen in small aliquots and stored at -80°C.

### Preparation of DNA templates for nucleosome reconstitution

All DNA templates were prepared by PCR amplification of the Widom 601 nucleosome positioning sequence using Phusion DNA polymerase and Phusion GC buffer (NEB). Primers were purchased from Eurofins Genomics. 147bp templates were prepared using a biotinylated forward primer (Bio-ctggagaatcccggtgccgaggc) and a reverse primer (acaggatgtatatatctgacacg). 175 bp templates were prepared by appending 14bp of linker DNA (underlined) to the primers above (forward primer Bio-<u>gacggacggacgga</u>ctggagaatcccggtgccgaggc, reverse primer <u>gacggacggacgga</u>acaggatgtatatatctgacacg). PCR reactions were diluted into 5 volumes of buffer A (10mM Tris pH 8) and DNA was purified using ion exchange chromatography on a RESOURCE Q column (Cytiva) using an elution gradient from 0-1M NaCl in 10mM Tris pH 8.

Positive fractions were pooled and DNA was precipitated by addition of 1/10^th^ volume of 3M NaOAc pH 5 and 3 volumes of ethanol and overnight incubation at -20°C. Precipitated DNA was pelleted by centrifugation at 5,000rpm at 4°C for 30 minutes. The pellet was dried, resuspended in TE (10mM Tris pH 8, 1mM EDTA), and stored at -20°C.

### Nucleosome reconstitution

Histones H2A, H2B, H3, and H4 were purchased from The Histone Source. Nucleosomes containing H3K18C or H3K18C-ub were reconstituted as described previously^36,38^. Briefly, histone octamers containing H3K18C or H3K18C-ub were formed by mixing individual histones in unfolding buffer (7M guanidine-HCl, 20mM Tris pH 7.5, 10mM DTT) and dialyzing into refolding buffer (10mM Tris pH 7.5, 2M NaCl, 1mM EDTA, 5mM BME). Octamers were mixed with DNA templates and nucleosomes were reconstituted using an 18 hour gradient salt dialysis from 2M to 250mM KCl. Nucleosome integrity was verified by native PAGE and SYBR safe staining.

### Nucleosomes for *in vitro* assays

Nucleosomes used in Figs. 1C, 2E, and 2F were produced according to the methods above. All other nucleosomes were purchased from Epicypher.

**Figure 1:**
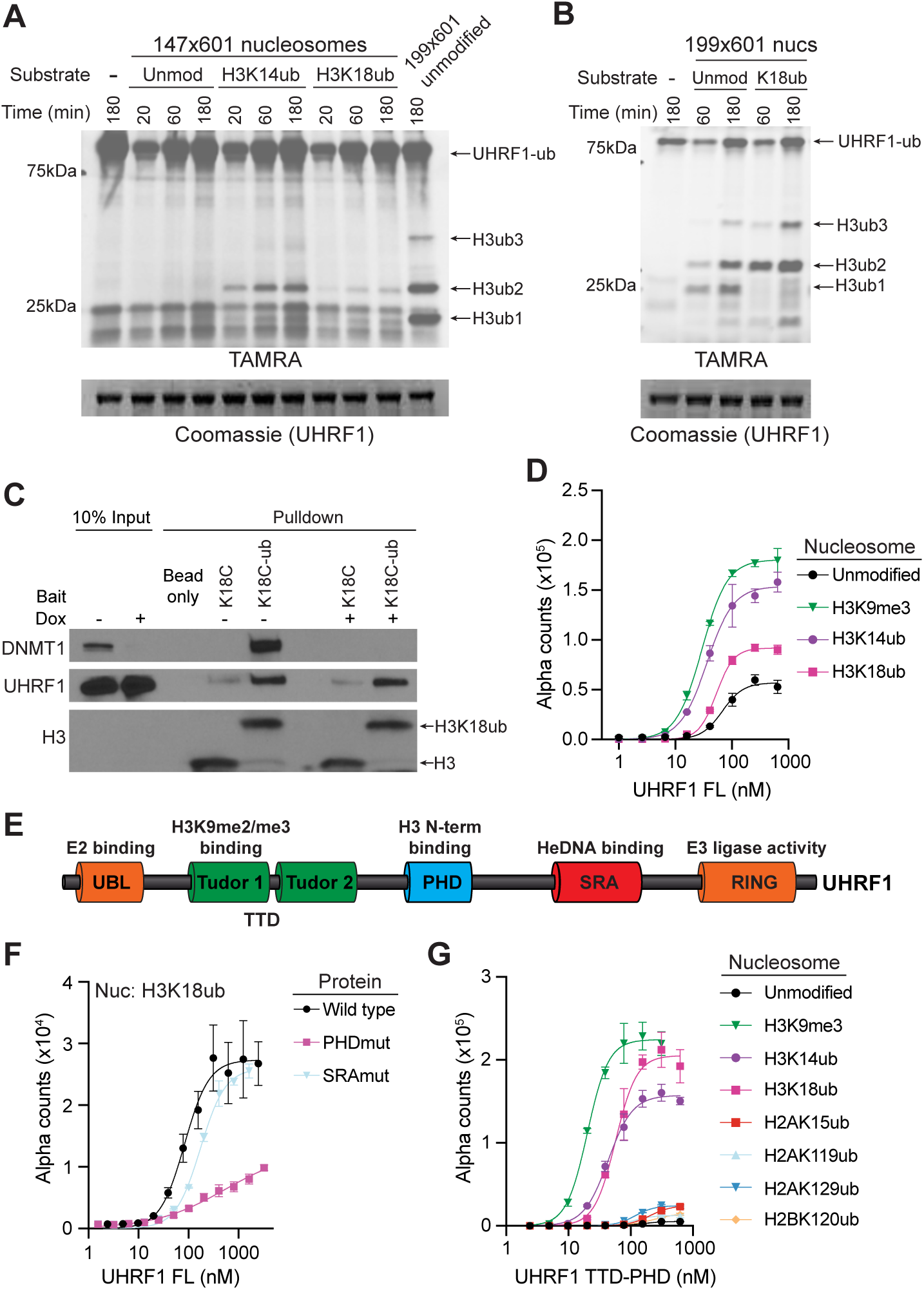
UHRF1 recognizes H3K18ub through its TTD-PHD domain. (**A-B**) *In vitro* ubiquitin ligase activity assays of recombinant UHRF1 using the indicated nucleosome substrates. TAMRA fluorescence shows ubiquitin and its products; Coomassie staining shows UHRF1. (**C**) Western blot analysis of proteins enriched from HCT116 dox-shDNMT1 nuclear extracts using biotinylated H3K18C or H3K18C-ub nucleosomes as bait. (**D**) AlphaScreen binding assays measuring the interaction between full-length UHRF1 and the indicated biotinylated nucleosomes. Error bars represent SD of triplicate reactions. (**E**) Domain architecture of UHRF1. (**F**) AlphaScreen binding assays measuring the interaction between full-length UHRF1 (wild type or indicated mutants) and the indicated biotinylated nucleosomes. Error bars represent SD of triplicate reactions. (**G**) AlphaScreen binding assays measuring the interaction between UHRF1 TTD-PHD and the indicated biotinylated nucleosomes. Error bars represent SD of triplicate reactions.

**Figure 2:**
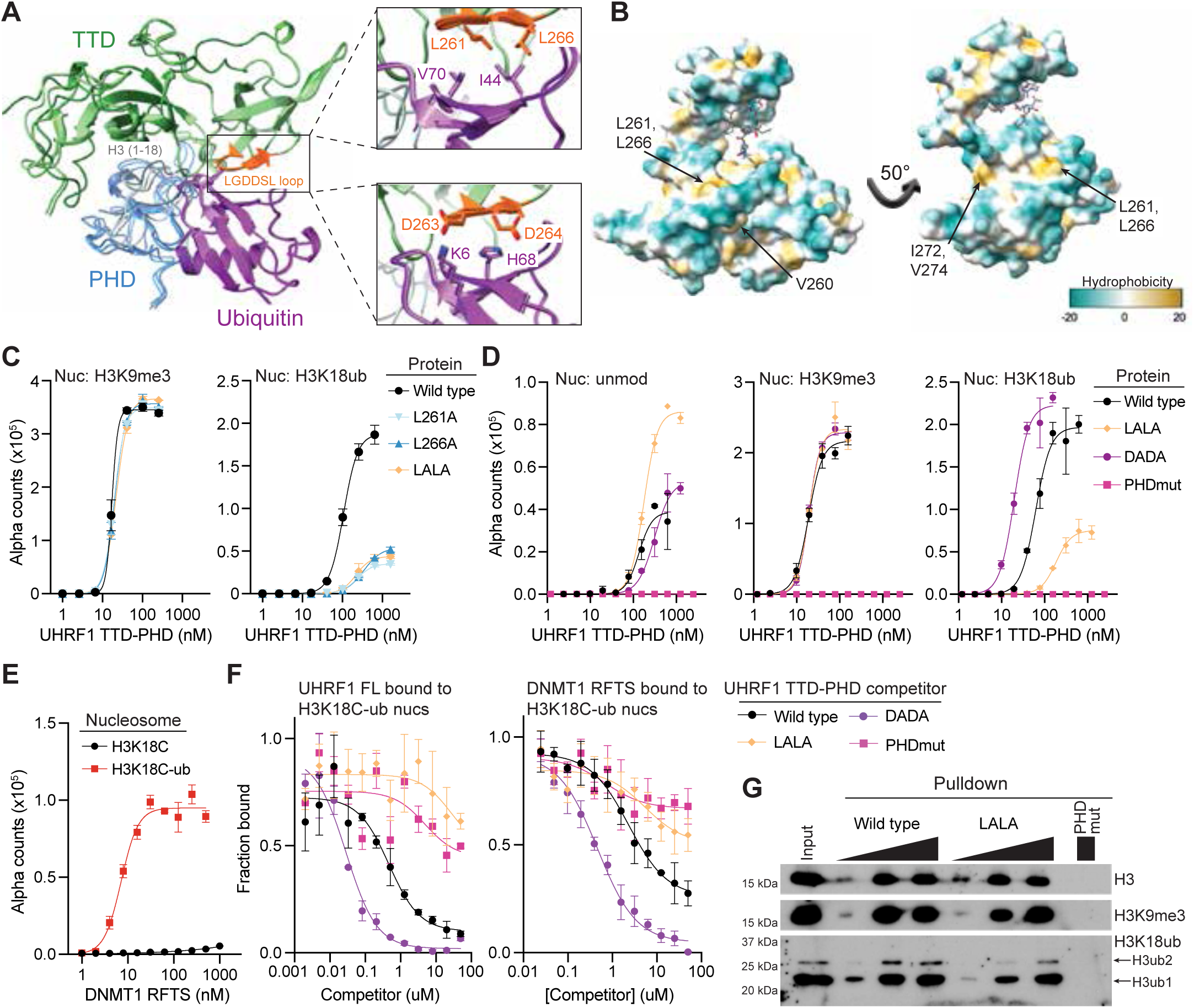
UHRF1 recognizes the ubiquitin hydrophobic patch through an LGDDSL loop on Tudor 2. (**A**) AlphaFold 2 predictions of a complex containing UHRF1 TTD-PHD, ubiquitin, and H3 (residues 1-18). Large box, overall predicted structure; small boxes, predicted interactions between residues in the LGDDSL loop of Tudor 2 and ubiquitin in the top three models. (**B**) Crystal structure of UHRF1 TTD-PHD in complex with an H3K9me3 peptide (PDB: 3ASK), with the protein surface colored by hydrophobicity. Hydrophobic surface residues mutated in (**C and S1A-B**) are indicated. (**C-D**) AlphaScreen binding assays measuring the interaction between wild-type or mutant UHRF1 TTD-PHD and the indicated biotinylated nucleosomes. Error bars represent SD of triplicate reactions. (**E**) AlphaScreen binding assays measuring the interaction between the DNMT1 RFTS and the indicated biotinylated nucleosomes. Error bars represent SD of triplicate measurements. (**F**) AlphaScreen competition assays measuring displacement of the interaction between H3K18C-ub nucleosomes and full length UHRF1 (left) or DNMT1 RFTS (right) in the presence of increasing concentrations of wild type or mutant UHRF1 TTD-PHD. Error bars represent SD of triplicate reactions. (**G**) Western blot analysis of HCT116 mononucleosomes enriched by pulldown with wild type or mutant UHRF1 TTD-PHD.

### AlphaFold structural predictions

*In silico* structural predictions of the complex of UHRF1 TTD-PHD, histone H3 aa1-18, and ubiquitin were performed using the official open-source local implementation of AlphaFold2 (v2.3.2)^39^ sourced directly from Google DeepMind. The multi-chain structures were predicted using the AlphaFold-Multimer (v3) neural network model^40^. Five distinct structural models were generated and ranked based on a weighted combination of the interface predicted local distance difference test (ipTM) and the global predicted TM-score (pTM). Confidence in the predicted backbone and interface orientations was verified using per-residue pLDDT values and Predicted Aligned Error (PAE) matrices. Alignment of the top three models is shown in **Fig. 2A**.

### AlphaScreen binding assays

Alpha screen assays with purified recombinant proteins were performed in alpha assay buffer (25mM HEPES pH 7.5, 250mM NaCl, 0.05% NP40) in 384 well light grey AlphaPlates (Revvity). Tagged recombinant protein was mixed with biotinylated nucleosome at 2x final concentration in a final volume of 10μL and incubated at room temperature for 30 minutes. Streptavidin donor beads and Ni NTA acceptor beads (for His-MBP-tagged proteins) or glutathione acceptor beads (for GST-tagged proteins) were diluted in alpha assay buffer and added to reactions for a final reaction volume of 20μL. Reactions were incubated at room temp for 30 minutes in the dark, then alpha counts were measured using a Biotek plate reader. In binding assays the final concentrations of nucleosome were 0.8nM for His-MBP-UHRF1 FL, 1.25nM for GST-TTD-PHD, and 5nM for His-MBP-RFTS. The final concentration of both donor and acceptor beads was 80μg/mL for His-MBP-tagged proteins and 40μg/mL for GST-tagged proteins. Data were plotted and fitted in GraphPad Prism using a specific binding with Hill slope model.

Alpha screen competition assays were performed as described above, except that three components were incubated together first (nucleosome, binding partner, and competitor), followed by addition of donor and acceptor beads. In competition assays the final concentrations were 0.8nM nucleosome + 750nM His-MBP-UHRF1, 5nM nucleosome + 80nM His-MBP-RFTS, and 2.5nM nucleosome + 125nM GST-TTD-PHD. The final concentration of both donor and acceptor beads was 80μg/mL for His-MBP-tagged proteins and 40μg/mL for GST-tagged proteins. Data were plotted and fitted in GraphPad Prism using an inhibitor vs. response (three parameters) model.

Alpha screen assays with E. coli lysates expressing GST-TTD-PHD were performed as above. The final concentration of nucleosome was 5nM and the final concentration of both donor and acceptor beads was 40μg/mL.

### *In vitro* UHRF1 ubiquitin ligase activity assays

Ubiquitin ligase activity assays were performed as previously described^34,41^ with minor modifications. Briefly, 20μL reactions were assembled in reaction buffer (50mM HEPES pH 7.5, 66mM NaCl, 2.5mM MgCl_2_, 2.5mM DTT) and included 50nM E1 (UBA1), 667nM E2 (UBCH5A), 1.5μM E3 (UHRF1), 1μM TAMRA-ubiquitin, and 8mM ATP. Reactions also contained either 100nM nucleosome substrate or 6.25μM hemimethylated 12bp DNA oligo and 1μM peptide substrate. Reactions were incubated at room temperature for the indicated time and stopped by addition of SDS sample buffer to 1x final and heating at 95C for 5 minutes. Reactions were separated by SDS-PAGE and gels were scanned for fluorescent visualization of TAMRA-ubiquitin, then stained with Coomassie blue.

### Mammalian cell culture

Cell lines used in this study included previously generated HCT116 cells (ATCC # CCL-247) containing a doxycycline (dox) inducible short hairpin RNA (shRNA) targeting the 3’ untranslated region of UHRF1 (HCT116 shUHRF1 c3)^35^ or DNMT1 (HCT116 shDNMT1 c3)^35^ and derivatives of the HCT116 shUHRF1 c3 cell line containing stably integrated shRNA-resistant UHRF1 transgenes. Cells were grown in McCoy’s 5A medium (Gibco 16600-082) supplemented with 10% FBS, 10U/mL penicillin/streptomycin (Gibco 15140122), and 8μg/mL blasticidin (to select for integrated transgenes) at 37°C and 5% CO_2_. Cells were passaged as necessary to maintain <80% confluency, typically every 2-3 days. For knockdown of endogenous UHRF1 or DNMT1, 10ng/mL doxycycline was added to the culture media.

### Preparation of HCT116 nuclear extracts

HCT116 shDNMT1 c3 cells were cultured as described above. For nuclear extract preparation, cells were washed with Dulbecco’s Phosphate Buffered Saline (DPBS) and removed from culture dishes using trypsin, washed using ice-cold PBS, and pellets were frozen in liquid nitrogen. Cell pellets were lysed in Buffer A (10mM HEPES-KOH, pH 7.9; 1.5mM KCl; 1.5mM MgCl_2_; 0.34M Sucrose; 0.2% NP-40; 10% glycerol) on ice for 10 minutes with gentle agitation every 5 minutes. Lysis reactions were centrifuged and the supernatant (cytosolic components) was removed leaving the pellet (intact nuclei). Nuclei were washed in Buffer B (10mM HEPES-KOH, pH 7.9; 1.5mM KCl; 1.5mM MgCl_2_; 150mM NaCl; 10% glycerol; 1mM DTT) and pelleted again. Nuclei were then resuspended and incubated in Buffer C (20mM HEPES-KOH, pH 7.9; 1.5mM MgCl_2_; 0.1% NP-40; 0.2mM EDTA; 350mM NaCl; 25% glycerol; 1mM DTT; 0.5mM PMSF) supplemented with RNase A/T1 to 2μg/mL with rotation at 15-30 RPM for 1 hour at 4°C. Nuclei were pelleted again and the supernatant (soluble nuclear fraction) was removed and quantified via BCA assay for input into pulldown reactions. Effective isolation of soluble nuclear proteins was verified via western blot.

### Preparation of mononucleosomes from HCT116 cells

Cell pellets were resuspended in lysis buffer (5mM PIPES pH 8, 85mM KCl, 1% NP40, 1x cOmplete protease inhibitors, 1mM PMSF, and 100μM PR619 deubiquitinase inhibitor) and incubated on ice for 20 minutes with occasional mixing to lyse. Samples were pelleted at 2000g at 4°C for 5 minutes and supernatant was discarded. The pellet (containing nuclei) was washed by resuspension in 1mL of MNase digestion buffer (50mM Tris pH 8, 1mM CaCl_2_, 300mM sucrose, 1mM PMSF, and 100μM PR619) and spun as before. The supernatant was discarded. To digest chromatin, the pellet was resuspended in 1mL of MNase digestion buffer supplemented with 50U of MNase and incubated at 37°C with shaking for 12.5 minutes.

Digestion was stopped by adding EDTA to 10mM final and transferring tubes to ice. To further digest and solubilize chromatin, samples were sonicated with a Covaris E220 evolution sonicator twice for 1 minute with the following parameters: peak power 140, duty factor 5, cycles/burst 200, temperature 4°C. Samples were centrifuged at 13,000rpm at 4°C for 10 minutes and soluble chromatin was used immediately for pulldowns or frozen at -80°C until use. To quantify chromatin, a 20μL input sample was saved and 80μL of elution buffer (50mM NaHCO_3_, 1% SDS), 6μL of 5M NaCl (300mM final), and 1μL of proteinase K (200μg/mL final) were added. Samples were incubated at 65°C overnight to reverse crosslinks and digest protein. The following day, RNA was digested by addition of 5μL of 2mg/mL RNaseA/T1 and incubation at 37°C for 30 minutes. DNA was purified using the Qiagen PCR purification kit and quantified using a Qubit fluorimeter.

### Pulldowns from mononucleosomes using recombinant GST-TTD-PHD

Recombinant GST-TTD-PHD was bound to magnetic glutathione agarose beads (40μL bed volume per reaction) in binding buffer (20mM Tris pH 8, 300mM NaCl, 1mM EDTA, 0.1% Triton X-100) at 4°C with rotation for at least 1 hour. Precomplexed beads were collected with a magnet and washed three times with binding buffer. Cellular mononucleosomes were supplemented with NaCl (300mM final) and Triton X-100 (0.1% final), then added to beads and allowed to bind at 4°C with rotation overnight. The following day the beads were collected with a magnet and washed three times with binding buffer, with 5 minutes of rotation at 4°C for each wash step. Beads were collected and resuspended in 1x SDS sample buffer (50mM Tris pH 6.8, 6% glycerol, 1% BME, 2% SDS, 0.02% bromophenol blue) and heated at 95C for 5 minutes to elute bound material and denature proteins. Samples were analyzed by SDS-PAGE and western blot.

### Pulldowns from nuclear extracts using biotinylated recombinant nucleosomes

Pulldown reactions were conducted as follows. Streptavidin Sepharose beads (Cytiva) were equilibrated in RB-low buffer (10 mM Tris-HCl, pH 7.5; 250 mM KCl; 1 mM EDTA; 0.1% NP-40) by washing and pelleting beads three times. After equilibration, 50 pmol of biotinylated nucleosome in RB-low buffer was added to equilibrated beads and rotated at 15-20 RPM at 4°C for pre-complexing. After pre-complexing the beads were pelleted and the supernatant removed. 50 μg of soluble nuclear lysates adjusted to 150 mM NaCl with adjustment buffer (20 mM HEPES-KOH, pH 7.9; 1.5 mM MgCl_2_; 0.1% NP-40; 0.2 mM EDTA; 10 mM NaCl; 25% glycerol; 1 mM DTT; 0.5 mM PMSF) was added to the pre-complexed beads and the reactions were rotated at 15-20 RPM at 4°C overnight. The next day, beads were pelleted and washed three times with pull-down buffer (20 mM HEPES-KOH, pH 7.9; 1.5 mM MgCl_2_; 0.1% NP-40; 0.2 mM EDTA; 150 mM NaCl; 25% glycerol; 1 mM DTT; 0.5 mM PMSF). Retained proteins and nucleosomes were then boiled in 1x SDS sample buffer (50mM Tris pH 6.8, 6% glycerol, 1% BME, 2% SDS, 0.02% bromophenol blue) and analyzed via SDS-PAGE and western blotting.

### Generation of HCT116 shUHRF1 cover cell lines

Retroviruses encoding transgene covers (empty vector, UHRF1 wild type, LALA, or DADA) were produced from the pMXs-IRES-blasticidin plasmid and used to transduce HCT116 shUHRF1 clone 3 cells as previously described^35^.

### Incucyte cell growth measurements

15,000 cells were plated per well in a 24-well plate in triplicate with or without 20ng/mL doxycycline as indicated. Cell confluency was measured on an Incucyte S3 Live-Cell Analysis System. 16 images were captured from each well at each timepoint.

### Genomic DNA preparation

For EPIC array analysis of DNA methylation, genomic DNA was isolated using phenol/chloroform extraction and ethanol precipitation. Cell pellets were resuspended in 2mL of TE (10mM Tris pH 8, 0.1mM EDTA) + 0.5% SDS + 1mg/mL proteinase K and incubated for 3 hours at 50°C to lyse cells and digest protein. DNA was extracted three times using first phenol, then phenol/chloroform/isoamyl alcohol (25:24:1), and finally chloroform/isoamyl alcohol (24:1). For each extraction, an equal volume of organic solvent was added and the sample was incubated on ice for 5 minutes, followed by centrifugation at 3,000rpm for 5 minutes at 4°C. The aqueous phase was recovered and used for the next extraction. After extraction, 10% volume of NaOAc pH 5 and 3 volumes of ethanol were added and the sample was incubated at -20°C overnight to precipitate DNA. Precipitated DNA was pelleted by centrifugation at 5,000rpm at 4°C for 30 minutes, washed three times with 70% ethanol, and allowed to dry. The pellet was resuspended in nuclease-free water and treated with 1mg/mL RNase A at 37°C for 1 hour. DNA was then precipitated and purified again as described and resuspended in nuclease-free water.

### EPIC arrays

Genomic DNA was quantified using the Qubit dsDNA high sensitivity quantification assay (Invitrogen) and submitted to the Van Andel Institute Genomics Core Facility for quality control, bisulfite conversion and DNA methylation quantification using Infinium MethylationEPIC BeadChIP v2 (Illumina) processed on an Illumina iScan system following the manufacturer’s instructions^42,43^.

### EPIC array analysis

All analyses were performed in the R statistical software (v4.4.2). The Bioconductor package ‘SeSAMe’ (v1.24.0) was used to process raw IDAT files, extract and normalize probe signal intensity values, and calculate β-values from the normalized probe signal intensity values^44–46^. CpG probes with a *P*-value > 0.05 in any one sample were excluded from analyses.

### Histone acid extraction

For extraction of histones from mammalian cells, cell pellets were resuspended in 1mL of hypotonic lysis buffer (10mM Tris pH 8, 1mM KCl, 1.5mM MgCl2, 1mM DTT, 1mM PMSF) and incubated with rotation at 4°C for 30 minutes to lyse cells by mechanical shearing. Samples were centrifuged at 10,000g for 10 minutes at 4°C to pellet nuclei. Supernatant was discarded and nuclei were washed with 1mL of hypotonic lysis buffer and pelleted as before. Supernatant was discarded and nuclei were resuspended in 400μL of 0.2M H_2_SO_4_ and incubated for 2-4 hours at 4°C with rotation. Samples were centrifuged at 16,000g at 4°C for 10 minutes and the supernatant was transferred to a new tube and mixed with 200μL TCA to precipitate histones. Samples were incubated at 4°C for 1 hour to overnight, then centrifuged at 16,000g at 4°C for 10 minutes to pellet histones. Supernatant was discarded and histones were washed 2-3 times with 500μL of acetone and pelleted at 3,400g at 4°C for 10 minutes. Pellets were air dried, then resuspended in RIPA buffer (20mM Tris pH 6.8, 1% SDS) by shaking at 37°C for 1 hour to overnight. Protein concentration was determined using the Pierce BCA Protein Assay Kit (Thermo Fisher Scientific).

### Western blots

Whole cell protein lysates were prepared by resuspending cell pellets in RIPA buffer (20mM Tris pH 6.8, 1% SDS) and sonicating to digest chromatin (10% amplitude, 0.2s on/0.8s off for 2s on total, repeated twice). Protein concentration was determined using the Pierce BCA Protein Assay Kit (Thermo Fisher Scientific).

Proteins (from whole cell lysates or acid extracted histones) were separated on SDS-PAGE gels and transferred to a PVDF membrane on a semi-dry transfer apparatus. Membranes were blocked with blocking buffer (PBST + 5% BSA) and probed with primary antibody in blocking buffer overnight at 4°C. Membranes were washed three times with PBST, probed with secondary antibody in blocking buffer for 1 hour, then washed three times with PBST. Membranes were probed with ECL substrate and imaged with film.

## RESULTS

### UHRF1 binds selectively to H3ub nucleosomes through its TTD-PHD domain

Although it is appreciated that UHRF1 can install multiple mono-ubiquitin marks on histone H3, it is unclear how successive ubiquitin additions are promoted and whether pre-installed H3 mono-ubiquitination influences UHRF1 catalytic activity. To study this, we compared the ubiquitin ligase activity of recombinant UHRF1 on semi-synthetic recombinant nucleosome substrates containing either unmodified H3 or site-specific mono-ubiquitination (H3ub1). While UHRF1 activity was undetectable toward 147×601 (linker-less) unmodified nucleosomes, the presence of H3K14ub, or to a lesser extent H3K18ub, stimulated the enzyme to write a second ubiquitin on H3 (H3ub2) (**Fig. 1A**). Consistent with our prior studies and the work of others^26,34^, UHRF1 enzymatic activity toward unmodified nucleosomes was enhanced when linker DNA was included, as shown by comparing its activity toward 147×601 nucleosomes versus 199×601 nucleosomes (26bp linkers) (**Fig. 1A-B**). The enhanced activity of UHRF1 toward H3K18ub nucleosomes compared to unmodified was further evident on substrates wrapped with linker DNA (**Fig. 1B**). As UHRF1 autoubiquitination was not increased in the presence of these H3ub1 substrates (**Fig. 1A-B**), these collective data suggested that the enhanced activity observed on H3ub1 nucleosomes was through superior targeting toward the H3 tail substrate.

We next sought to determine whether H3ub1 nucleosomes could serve as bait to enrich for endogenous UHRF1 from nuclear extracts. To test this, we reconstituted mononucleosomes wrapped with 175bp of biotinylated DNA containing H3K18C with a dichloroacetone linkage to ubiquitin G76C, which mimics a native ubiquitin linkage but is resistant to hydrolysis by deubiquitinases^47^. Nuclear extracts were prepared from HCT116 colon cancer cells expressing doxycycline-inducible shRNA to knock down DNMT1, a key binding partner of UHRF1 and the only reported high-affinity reader of H3ub^32^. The preferential interaction of DNMT1 with H3K18C-ub nucleosomes over their unmodified controls was verified in this assay system (**Fig. 1C**). Like DNMT1, UHRF1 was also enriched preferentially with ubiquitinated nucleosomes.

Importantly, this enrichment was maintained upon DNMT1 depletion, further supporting the hypothesis that UHRF1 directly interacts with H3ub1 nucleosomes.

To directly determine the effect of H3ub1 on UHRF1 interaction with nucleosomes, we next performed AlphaScreen proximity-based binding assays between recombinant full-length UHRF1 (UHRF1 FL) and linker-less semi-synthetic nucleosomes. Consistent with the reported activities of the TTD-PHD histone binding unit, concentration-dependent interaction of UHRF1 with unmodified nucleosomes was measurable, and this interaction was enhanced by the addition of H3K9me3 (**Fig. 1D**). Notably, both H3K14ub and H3K18ub enhanced the interaction of UHRF1 FL with nucleosomes, demonstrating that UHRF1 binding to nucleosomes is increased by pre-marked H3K14ub and H3K18ub.

We next sought to characterize the specificity and selectivity of this newly identified interaction. The appreciated nucleosome binding surfaces of UHRF1 are its DNA-binding SRA domain^23–25,28^ and its histone-binding TTD-PHD^18,19,21^ (**Fig. 1E**). The PHD anchors UHRF1 to the unmodified N-terminus of H3 and this interaction is enhanced through binding of the first Tudor in the TTD to H3K9me3 through an aromatic cage. While a point mutation in the SRA that perturbs its DNA binding activity (G448D)^28^ had minimal effect on UHRF1 FL interaction with H3K18ub nucleosomes, point mutations in the UHRF1 PHD that disrupt its interaction with the H3 tail (D334A/E335A)^19,21^ strongly impacted binding (**Fig. 1F**). These data suggest that the binding specificity of UHRF1 to H3K18ub nucleosomes is harbored in its TTD-PHD and, like H3K9me3, is dependent on multivalent anchoring through its PHD. Consistent with this hypothesis, the binding preference for H3K9me3, H3K14ub, and H3K18ub over unmodified nucleosomes was retained in the isolated TTD-PHD (**Fig. 1G**). Moreover, nucleosomes bearing other known histone ubiquitination marks (H2AK15ub, H2AK119ub, H2AK129ub, and H2BK120ub) were inferior binding partners for the TTD-PHD compared to nucleosomes ubiquitinated on H3, demonstrating that this newly identified reading activity has selectivity for ubiquitin sites on the H3 tail over other sites tested (**Fig. 1G**). Collectively, these data demonstrate that UHRF1 directly and selectively recognizes its mono-ubiquitinated H3 products through its TTD-PHD and suggest that this interaction enhances substrate engagement to promote successive ubiquitin deposition.

### An LGDDSL loop within Tudor 2 mediates ubiquitin recognition by the UHRF1 TTD-PHD

Having established that the UHRF1 TTD-PHD selectively binds H3 mono-ubiquitination, we next sought to define the region of this histone binding unit responsible for ubiquitin recognition. To identify candidate residues mediating ubiquitin engagement, we built an AlphaFold2 model of the UHRF1 TTD-PHD in complex with H3 (residues 1-18) and ubiquitin. Because AlphaFold2 does not accept ubiquitin as a post-translational modification (PTM), the interaction was modeled using three separate protein/peptide chains. The three top-ranked models predicted the reported interaction between the PHD and the H3 N-terminus from x-ray crystallography studies^21^ with some degree of variability that is consistent with our prior molecular dynamics simulations19. Notably, these models converged on predicted interactions between ubiquitin and a surface-exposed loop within Tudor 2 (residues 261-266; LGDDSL loop) (**Fig. 2A**). Specifically, the models predicted packing of UHRF1 L261 and L266 against I44 and V70 of the hydrophobic patch on ubiquitin and potential salt bridge interactions between UHRF1 D263/D264 and ubiquitin K6/H68.

Because many of the reported ubiquitin binding protein interactions involve a hydrophobic patch^48^, we next tested whether hydrophobic residues within this Tudor 2 surface contribute to H3ub1 recognition. Combinations of alanine point mutations were introduced into L261, L266, and adjacent hydrophobic residues V260, I272, and V274 (**Fig. 2B**), and these wild-type (WT) and mutant forms of recombinant UHRF1 TTD-PHD were tested in AlphaScreen assays for binding to semi-synthetic modified nucleosomes. While each of these mutations preserved binding of crude and purified TTD-PHD to H3K9me3-containing nucleosomes, L261A and L266A disrupted binding to H3K14ub- and H3K18ub-containing nucleosomes (**Fig. 2C, S1A-B**). These data indicate that L261 and L266 are specifically required for UHRF1 recognition of ubiquitinated H3 and that this interaction is separable from recognition of H3K9me3.

We next asked whether the salt bridges predicted by AlphaFold2 involving UHRF1 D263 and D264 were important for recognition of ubiquitin. Like the identified leucine mutations that disrupt H3ub1 binding, UHRF1 TTD-PHD D263A/D264A (DADA) bound unmodified and H3K9me3-containing nucleosomes similarly to WT (**Fig. 2D**). However, DADA enhanced UHRF1 binding to H3K14ub- and H3K18ub-containing nucleosomes (**Fig. 2D, S1C**). These data indicate that while the LGDDSL loop defines the ubiquitin interaction surface, the precise atomic contacts predicted by AlphaFold2 modeling do not fully capture the details of the interaction.

While AlphaScreen binding assays are a powerful method for detecting molecular interactions, signal intensity is influenced by proximity geometry, bead avidity, and saturation, limiting its utility for making quantitative measurements of binding affinity. In contrast, competition-format AlphaScreen measures displacement of a defined interaction and therefore provides a more reliable estimate of relative binding affinities in the nanomolar to micromolar range. To quantitatively assess the effects of these point mutations on the affinity of the UHRF1 TTD-PHD for H3ub1, we performed competition AlphaScreen assays to displace the RFTS domain of DNMT1 (the known H3ub1 reader; **Fig. 2E**) or full length UHRF1 (**Fig. 1F**). The WT TTD-PHD effectively competed with both UHRF1 FL and DNMT1 RFTS for binding to H3K18C-ub-containing nucleosomes, whereas competition with LALA- and PHD-mutant forms was impaired (**Fig. 2F, S1D**). In contrast, competition with the DADA-mutant UHRF1 TTD-PHD was enhanced compared to WT, consistent with its apparent enhanced affinity in the binding assay format.

We next asked whether this interaction occurs in the context of cellular chromatin. Mononucleosomes isolated from HCT116 colon cancer cells were used as bait for pulldowns with recombinant GST-TTD-PHD. Consistent with our *in vitro* observations, wild type GST-TTD-PHD enriched nucleosomes marked by H3K9me3 and H3K18ub, whereas a PHD-mutant protein failed to enrich for either modification (**Fig. 2G**), demonstrating that recognition of both modified nucleosome populations requires PHD-mediated H3 tail anchoring. Compared to WT, the LALA-mutant TTD-PHD was impaired for enrichment of H3K18ub nucleosomes, while binding to unmodified H3 or H3K9me3 was unaffected. These data confirm that the LALA mutation specifically disrupts the interaction between the TTD-PHD and H3ub on cellular chromatin without perturbing the interaction with H3K9me3. Collectively, these data identify the LGDDSL loop within UHRF1 Tudor 2 as a previously unrecognized ubiquitin binding surface that enhances PHD-anchored recognition of mono-ubiquitinated H3 and is functionally separable from H3K9me3 recognition by Tudor 1.

### UHRF1 recognition of H3ub promotes multi-mono-ubiquitination

We next sought to investigate the contribution of this newly identified ubiquitin reading function to UHRF1 enzymatic activity. Using 199×601 nucleosomes containing unmodified H3, we first compared the catalytic activities of recombinant WT UHRF1 with DNA-binding and ubiquitin-binding mutants. Whereas WT UHRF1 catalyzed time-dependent accumulation of H3ub1, H3ub2, and H3ub3 products, a G448D mutation that disrupts SRA-mediated DNA binding^28^ strongly inhibited UHRF1 E3 ligase activity toward unmodified 199×601 nucleosomes (**Fig. 3A**). These data are consistent with the appreciated role for linker DNA enhancing UHRF1 enzymatic activity toward H3^34^. In contrast, the ubiquitin binding-deficient LALA mutant retained the ability to catalyze H3ub1 but was markedly impaired in generating H3ub2 and H3ub3 products.

**Figure 3:**
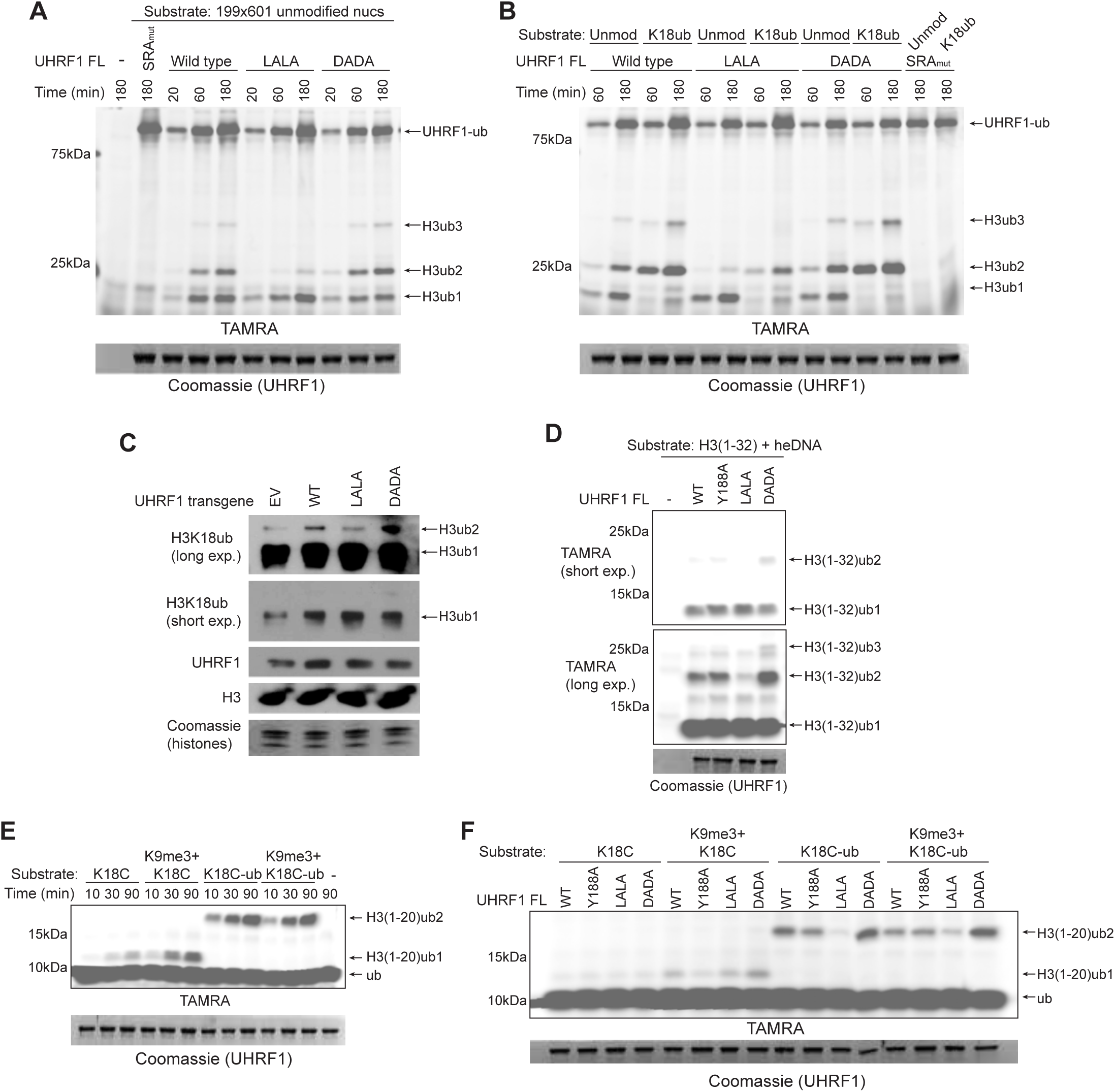
UHRF1 recognition of H3ub promotes multi-mono-ub *in vitro* and in cells. (**A-B**) *In vitro* ubiquitin ligase activity assays of recombinant UHRF1 using the indicated nucleosome substrates. TAMRA fluorescence shows ubiquitin and its products; Coomassie staining shows UHRF1. (**C**) Proteins from HCT116 shUHRF1 c3 cells expressing the indicated UHRF1 transgene covers analyzed by western blots and Coomassie staining as indicated. (**D-F**) *In vitro* ubiquitin ligase activity assays of wild-type or mutant UHRF1 using the indicated H3 peptide substrates.

Notably, autoubiquitination of UHRF1 was comparable between WT, SRA-mutant, and LALA-mutant forms of the enzyme, indicating that these mutations do not compromise intrinsic catalytic activity but rather specifically affect mono-ubiquitinated H3 substrate recognition. The ubiquitin binding-competent UHRF1 DADA mutant retained the ability to write multi-mono-Ub on H3.

These observations support a model in which ubiquitin recognition by UHRF1 promotes successive H3 ubiquitination. We therefore hypothesized that nucleosomes bearing a pre-installed H3ub1 would serve as superior substrates for WT UHRF1 but not for the ubiquitin binding-deficient LALA mutant. Consistent with this hypothesis, H3K18ub-containing nucleosomes were more efficiently ubiquitinated by WT UHRF1 and the DADA mutant than unmodified nucleosomes (**Fig. 3B**). In contrast, the LALA mutant showed reduced activity toward H3K18ub-containing nucleosomes, both relative to unmodified nucleosomes and to WT UHRF1 and DADA on the same H3K18ub substrate.

We next asked whether this ubiquitin-dependent read-write activity similarly promotes H3 multi-mono-ubiquitination in cells. To test this, we leveraged a clonally expanded HCT116 cell line engineered to express a doxycycline-inducible shRNA targeting the 3’ untranslated region of endogenous UHRF1^35^. We introduced shRNA-resistant transgenes encoding an empty vector (EV) or WT, LALA-mutant, or DADA-mutant UHRF1. After confirming UHRF1 overexpression in these cell lines (**Fig. 3C**), acid-extracted histones were analyzed for H3K18ub by western blotting. As expected, cells overexpressing WT UHRF1 had higher levels of both H3ub1 and H3ub2 than EV cells (**Fig. 3C**). Overexpression of LALA and DADA mutants increased H3ub1 to a similar extent as WT UHRF1. However, consistent with our *in vitro* assays, LALA-mutant UHRF1 was impaired for writing H3ub2, whereas DADA-mutant UHRF1 supported enhanced H3ub2 accumulation. Collectively, these results demonstrate that ubiquitin recognition is dispensable for H3 mono-ubiquitination, but necessary to promote successive multi-mono-ubiquitination of H3 *in vitro* and in cells.

### Tudor 1 and Tudor 2 independently stimulate UHRF1 E3 ligase activity

We next asked whether the H3K9me3 and H3ub reading functions of UHRF1 Tudor 1 and Tudor 2, respectively, function together or independently to regulate UHRF1 enzymatic activity. To distinguish between these possibilities, we compared the catalytic activity of WT UHRF1 with the LALA and DADA ubiquitin binding Tudor 2 mutants and Y188A, a previously characterized Tudor 1 mutant that disrupts recognition of H3K9me3^20^. Comparative activities were first analyzed with H3 peptide substrates (amino acids 1-32) in the presence of hemi-methylated DNA, which was included to accelerate UHRF1 enzymatic activity toward peptide substrates^28^. Consistent with our activity assays with nucleosome substrates, WT and mutant UHRF1 proteins exhibited comparable H3ub1 writing activity, whereas the LALA mutant was impaired for H3ub2 and H3ub3 writing activity and the DADA mutant showed enhanced H3ub2 and H3ub3 writing activity **(Fig. 3D).** The Y188A mutant was indistinguishable from WT UHRF1 in writing multi-mono-ub on these peptides that lacked H3K9me3, indicating that the absence of H3K9me3 binding by Tudor 1 does not impair Tudor 2-dependent recognition and successive ubiquitination of H3ub.

To directly test whether Tudor 1 and Tudor 2 cooperate when their respective binding moieties are present on the same H3 tail, we next generated a panel of H3 peptides (amino acids 1-20) with K14C or K18C, either unmodified or conjugated to ubiquitin G76C, together with unmodified or tri-methylated H3K9 **(Fig. S2A)**. WT UHRF1 ubiquitinated both K14C- and K18C-containing peptides but was more active toward K14C substrates (adding ub to K18) than toward K18C substrates (adding ub to K14) **(Fig. 3E, S2B)**. This *in vitro* substrate preference is consistent with prior data showing that H3K18 is a major site of UHRF1-mediated ubiquitination in cells^28^. In agreement with our nucleosome-based activity assays, peptides with pre-installed ubiquitin at either K14C or K18C enhanced subsequent ubiquitination by UHRF1 relative to the corresponding unmodified peptide **(Fig. 3E, S2B)**. H3K9me3 similarly enhanced UHRF1 activity toward both K14C and K18C substrates. However, combining H3K9me3 with K14C-ub or K18C-ub on the same peptide did not further stimulate ubiquitination beyond that observed with either modification alone, suggesting that Tudor 1 and Tudor 2 reading activities function independently **(Fig. 3E, S2B)**.

We next used Tudor 1 (Y188A) and Tudor 2 (LALA and DADA) mutants to determine whether the stimulatory effects of H3K9me3 and H3ub could be functionally uncoupled. The Y188A mutation abolished the increased activity of UHRF1 on H3K9me3 peptides while preserving the stimulatory effect of pre-installed ubiquitin at K14C or K18C **(Fig 3F, S2C)**. Conversely, LALA and DADA mutations abrogated and enhanced, respectively, the stimulatory effect of pre-installed ubiquitin without altering the response to H3K9me3 **(Fig 3F, S2C)**. Together, these data show Tudor 1 recognition of H3K9me3 and Tudor 2 recognition of H3ub provide independent, functionally separable inputs that stimulate UHRF1 E3 ligase activity.

### UHRF1 ubiquitin reading activity reinforces DNA methylation maintenance at CpG-sparse genomic regions

Finally, we asked whether the newly identified ubiquitin reading activity of UHRF1 contributes to DNA methylation maintenance. To address this, we employed a previously established genetic complementation strategy in which endogenous UHRF1 is depleted by doxycycline (dox)-inducible shRNA and replaced with shRNA-resistant WT or mutant UHRF1 transgenes^35,49^ (**Fig. 4A**). DNA methylation was measured using the Infinium (Illumina) MethylationEPIC BeadChip (EPIC array), which interrogates approximately 900,000 individual CpGs distributed throughout the genome.

**Figure 4:**
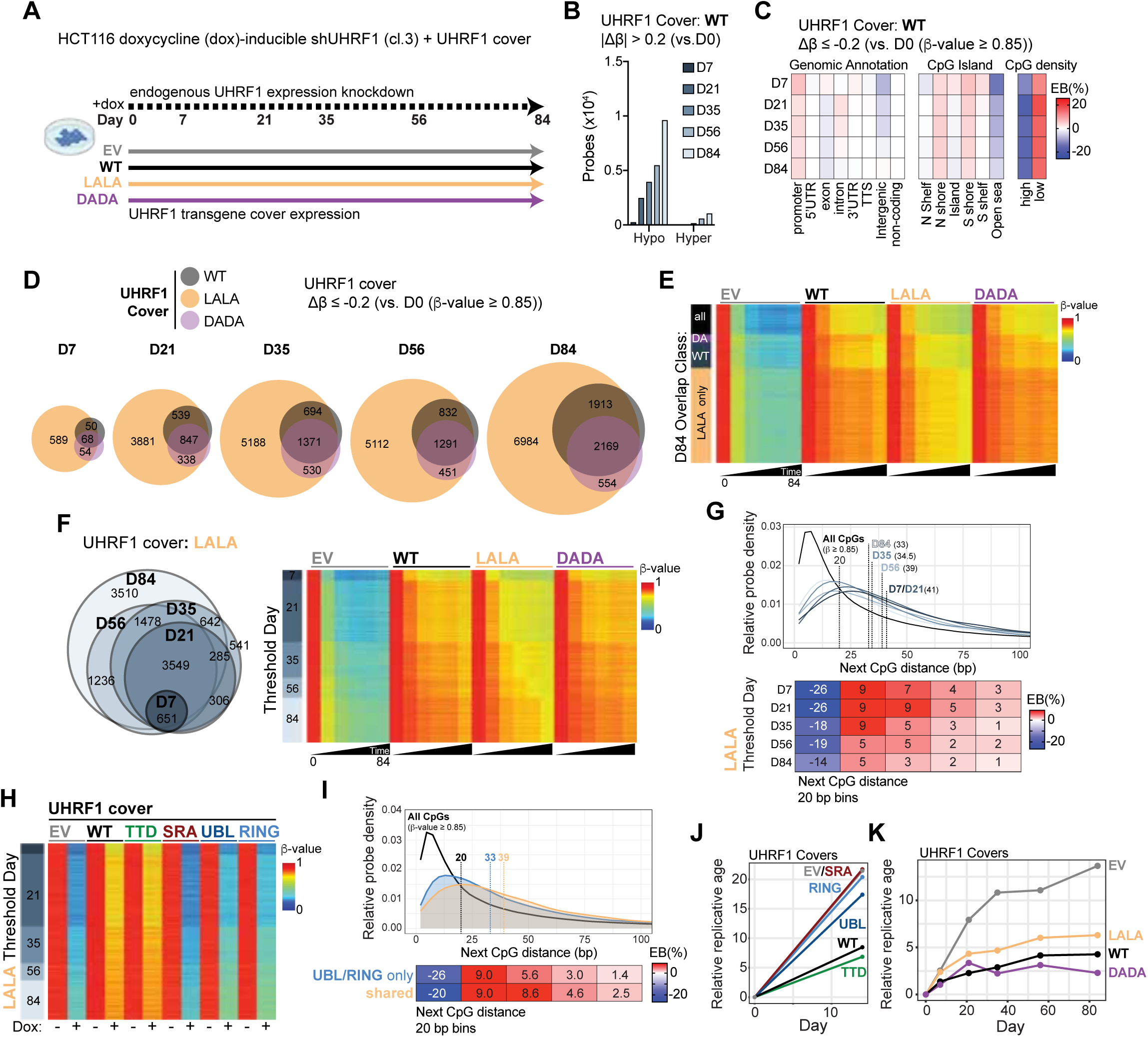
UHRF1 recognition of H3ub supports DNA methylation maintenance. (**A**) Experimental design for HCT116 cells expressing doxycycline-inducible shRNA targeting endogenous UHRF1 together with empty vector (EV) or shRNA-resistant wild-type (WT), LALA, or DADA UHRF1 transgenes. Following induction of endogenous UHRF1 knockdown, cells were maintained for 84 days and collected at the indicated time points for EPIC array analysis. (**B**) Number of probes exhibiting DNA methylation changes (|Δβ| > 0.2 relative to day 0) at the indicated time points in cells expressing WT UHRF1 cover, separated into hypomethylated and hypermethylated probes. (**C**) Enrichment analysis of genomic annotations, CpG island context, and CpG density among hypomethylated probes (Δβ ≤ −0.2) relative to highly methylated day 0 probes (β-value ≥ 0.85) in cells expressing WT UHRF1 cover. Color represents enrichment bias (EB). (**D**) Overlap of hypomethylated probes (Δβ ≤ −0.2 relative to day 0 β-value ≥ 0.85) identified at the indicated time points in cells expressing WT, LALA, or DADA UHRF1 covers. (**E**) DNA methylation β-values across the 84-day time course for probes grouped by their day-84 overlap classification among WT, LALA, and DADA UHRF1 cover conditions (from **D**). Heatmaps show methylation dynamics in EV, WT, LALA, and DADA cells over all collected time points. (**F**) Temporal accumulation of hypomethylated probes in LALA-expressing cells. The Venn diagram shows overlap among probes reaching the hypomethylation threshold (Δβ ≤ −0.2) at the indicated time points; heatmaps show DNA methylation dynamics for probes grouped by the first day at which the hypomethylation threshold was reached. (**G**) Top: density distribution of distances to the next CpG for probes reaching the hypomethylation threshold at the indicated time points in LALA-expressing cells. Density curves are compared with all highly methylated CpGs at baseline (β-value ≥ 0.85). Bottom: heatmap of enrichment bias for next CpG distance binned at 20 bp intervals from 2-100 bp. (**H**) DNA methylation dynamics of LALA threshold probes in cells expressing EV or UHRF1 WT, TTD, SRA, UBL, or RING transgene covers (from reference^35^), with or without doxycycline-induced depletion of endogenous UHRF1. (**I**) Top: density distribution of distances to the next CpG for hypomethylated CpG probes UBL/RING-only and shared probe populations identified in **Fig. S3G**, compared with all highly methylated CpGs at baseline. Bottom: heatmap of enrichment bias for next CpG distance binned at 20 bp intervals from 2-100 bp. (**J**) Relative replicative age (RepliTali) of cells expressing the indicated UHRF1 transgene covers in the presence or absence of doxycycline-mediated endogenous UHRF1 depletion (from reference^35^). (**K**) Relative replicative age across the 84-day time course in cells expressing EV, WT, LALA, or DADA UHRF1 transgene covers.

As expected, depletion of endogenous UHRF1 in cells carrying an empty vector (EV) transgene resulted in substantial DNA hypomethylation after 7 days of dox treatment (**Fig. S3A**). In contrast, WT-, LALA-, and DADA-expressing cover lines all maintained DNA methylation during this interval, with only a modest reduction in DNA methylation observed in cells expressing the ubiquitin reading-defective LALA mutant (**Fig. S3A**). This phenotype contrasts with previously characterized UHRF1 reader (PHD and SRA) and writer (UBL and RING) mutants, which induce substantial DNA hypomethylation within 7-14 days^35,49^. These observations suggest that ubiquitin recognition is largely dispensable for global maintenance methylation over short timescales.

We previously showed that UHRF1 E3 ubiquitin ligase activity is particularly important for maintaining DNA methylation at late-replicating, CpG-sparse regions that are susceptible to methylation loss during oncogenesis, aging, and serial passaging^11,12,35^. We therefore hypothesized that impairment of ubiquitin recognition might selectively compromise DNA methylation maintenance at these vulnerable loci over repeated cell divisions. To test this hypothesis, we monitored DNA methylation dynamics in WT-, LALA-, and DADA-expressing cover cell lines over 84 days in culture (**Fig. 4A**). Importantly, all cell lines proliferated at comparable rates **(Fig. S3B)** and exhibited nearly identical DNA methylomes at the beginning of the experiment (D0) **(Fig. S3C)**.

We first defined CpGs in WT UHRF1-expressing cells that undergo progressive DNA methylation loss during serial passaging. DNA methylation changes were quantified at each timepoint relative to day 0 using a |Δβ|-value cutoff of ≥ 0.2. Serial passaging of cells expressing WT UHRF1 resulted in progressive DNA hypomethylation over time (**Fig. 4B**), and these sites were strongly enriched among CpG-sparse regions **(Fig. 4C**). Based on their progressive loss of methylation through repeated cell divisions, we refer to these sites as replication-vulnerable CpGs.

We next examined the impact of disrupted UHRF1 ubiquitin recognition on DNA methylation dynamics at these replication-vulnerable CpG sites. Notably, cells expressing the LALA mutant accumulated substantially more hypomethylated CpGs than cells expressing WT or DADA UHRF1 at every timepoint examined **(Fig. 4D-E).** By day 84, the number of hypomethylated CpGs in LALA-expressing cells exceeded that observed in WT- and DADA-expressing cells by more than 3-fold. Importantly, the majority of replication-vulnerable CpGs identified in WT-expressing cells were contained within this expanded set of hypomethylated loci (**Fig. 4D**). Moreover, the additional CpGs that became susceptible to methylation loss in LALA-expressing cells were similarly enriched among CpG-sparse regions (**Fig. S3D**). These findings show that disruption of UHRF1 ubiquitin recognition expands the population of CpGs susceptible to replication-associated DNA methylation erosion.

To compare kinetics of methylation erosion among the UHRF1 cover lines, we next grouped hypomethylated CpGs according to the timepoint (day) at which they reached a Δβ ≤ -0.2 threshold during serial passaging relative to CpGs with baseline β-value ≥ 0.85 at day 0 (D0). Among replication-vulnerable CpGs identified in WT-expressing cells, methylation loss occurred earlier and was more extensive in LALA-expressing cells, whereas methylation erosion was modestly attenuated in cells expressing the DADA mutant **(Fig. S3E-F).** Conversely, WT- and DADA-expressing cells largely maintained DNA methylation at the additional replication-vulnerable CpGs that became progressively hypomethylated in LALA-expressing cells (**Fig. 4F, S3F**). Notably, CpGs that crossed the hypomethylation threshold earliest in LALA-expressing cells (D7-D21) were strongly enriched for CpG-sparse regions **(Fig. 4G**). Together, these findings further demonstrate that UHRF1 ubiquitin recognition reinforces DNA methylation maintenance at replication-vulnerable CpGs.

An important prediction of this model is that CpGs rendered vulnerable by loss of UHRF1 ubiquitin recognition should also depend on UHRF1-mediated histone ubiquitination. To test this hypothesis, we integrated our previously published EPIC array datasets generated from cells expressing catalytically impaired UHRF1 UBL and RING mutants^35^. Consistent with this hypothesis, both the replication-vulnerable CpGs identified in WT-expressing cells and the expanded set revealed in LALA-expressing cells showed strong dependence on UHRF1 E3 ligase activity **(Fig. 4H)**. Thus, the genomic regions most sensitive to disruption of ubiquitin recognition are the same regions that require UHRF1-dependent histone ubiquitination for faithful maintenance methylation. However, only a subset of CpGs dependent on UHRF1 E3 ligase activity became replication-vulnerable upon disruption of ubiquitin recognition **(Fig. S3G**). These overlapping loci were further biased toward CpG-sparse regions compared to the full set of CpGs dependent on UHRF1 E3 ligase activity (**Fig. 4I).** Collectively, these findings functionally connect UHRF1 recognition of its H3ub products with its ubiquitin writing activity and identify CpG-sparse regions as particularly dependent on this read-write mechanism for DNA methylation maintenance.

DNA methylation loss within late replicating, CpG-sparse regions of the genome is a hallmark of both oncogenesis and aging^11,12,35^. Indeed, previous work by Endicott *et al.* leveraged this progressive replication-associated methylation loss to develop RepliTali (<u>Repli</u>cation <u>T</u>imes <u>A</u>ccumulated in <u>Li</u>fetime), an EPIC array-based metric that estimates the relative replicative history of cells^12^. We therefore asked whether disruption of UHRF1 ubiquitin reading and writing activities altered this replication-associated methylation signature. Disruption of ubiquitin writing through UBL or RING mutations^35^ increased predicted replicative age relative to UHRF1 WT-expressing cells (**Fig. 4J**). Similarly, LALA-expressing cells exhibited increased predicted replicative age relative to WT- and DADA-expressing cells as early as day 7, and this difference persisted throughout the timecourse **(Fig. 4K)**. Collectively, these findings demonstrate that UHRF1 ubiquitin reading activity protects against DNA methylation erosion at replication-vulnerable CpGs, linking the feed-forward read-write mechanism that promotes H3 multi-mono-ubiquitination to the reinforcement of DNA methylation maintenance in genomic regions that are particularly susceptible to progressive hypomethylation during repeated cell divisions.

## DISCUSSION

In this study, we identify a previously unrecognized ubiquitin reading activity in UHRF1 that promotes successive ubiquitination of histone H3 and reinforces DNA methylation inheritance. UHRF1 directly binds its mono-ubiquitinated H3 products through a surface-exposed LGDDSL loop in Tudor 2, with binding anchored by PHD engagement of the H3 N-terminus. Disruption of this interaction selectively impairs deposition of additional ubiquitin marks without preventing initial H3 mono-ubiquitination, whereas enhancing H3ub recognition promotes H3 multi-mono-ubiquitination. Importantly, these effects are observed both *in vitro* and in cells and translate into altered DNA methylation maintenance during repeated cell divisions. Together, our findings support a feed-forward read-write mechanism in which recognition of an H3ub product promotes its use as a substate for further ubiquitin deposition, thereby reinforcing DNA methylation inheritance at replication-vulnerable CpGs in the genome.

Tudor domains are found in many epigenetic regulators and are well-characterized as readers of lysine and arginine methylation that couple these modifications to downstream chromatin functions^50,51^. To our knowledge, the LGDDSL loop identified here represents the first example of a Tudor domain surface contributing to ubiquitin recognition. Of the numerous ubiquitin binding proteins identified to date, most utilize relatively low affinity interactions with ubiquitin^48^. Consistent with this, UHRF1 did not detectably bind to H3ub when PHD-mediated anchoring to the H3 N-terminus was disrupted, suggesting that proximity created by the PHD-H3 interaction enables productive engagement of ubiquitin by Tudor 2. This organization parallels recognition of H3K9me2/me3 by Tudor 1, which is similarly supported by PHD engagement of the H3 N-terminus. However, our findings indicate that the two Tudor domains provide functionally separable inputs to UHRF1. Disruption of the Tudor 2 ubiquitin binding surface did not impair H3K9me3 recognition, and disruption of the Tudor 1 H3K9me3 binding aromatic cage did not impair H3ub-dependent activity. Moreover, H3K9me3 and H3ub independently stimulated UHRF1 activity without evidence of cooperativity when present on the same H3 tail. Thus, the TTD-PHD functions as a multivalent histone recognition module in which a common PHD anchor supports distinct PTM reading activities in Tudor 1 and Tudor 2. More broadly, other chromatin readers that lack annotated ubiquitin binding domains may similarly engage ubiquitin through low-affinity surfaces that become functional only when coupled with higher affinity interactions with neighboring features on the same substrate.

Read-write mechanisms in which recognition of pre-existing histone ubiquitination directs subsequent modifications are well-documented in chromatin biology^52,53^. This has primarily been studied as a mechanism to regulate histone methylation^54–59^, and the H3K14ub and H3K18ub modifications studied here have been reported to stimulate H3K9 methylation as a mechanism to support heterochromatin maintenance^49,60,61^. The ubiquitin read-write mechanism described here is conceptually distinct and provides an explanation for how UHRF1 generates multi-mono-ubiquitinated H3 after initially encountering an unmodified substrate. Although multi-mono-ubiquitination has been described on a number of proteins with diverse functional consequences^62–66^, this is the first study to demonstrate that mono-ubiquitin recognition can directly promote ubiquitin writing at a proximal site to generate multi-mono-ubiquitination. Analogous product recognition mechanisms are likely to exist for other E3 ligases that generate multi-mono-ubiquitinated substrates.

Reciprocal phenotypes of the LALA and DADA mutants further support a direct relationship between H3ub recognition and successive ubiquitin deposition. However, structural studies will be required to define the atomic basis of ubiquitin engagement by the LGDDSL loop and to understand why substitution of D263 and D264 enhances rather than disrupts this interaction. The separation-of-function LALA and DADA mutants also provide insight into why UHRF1 deposits multiple mono-ubiquitin marks on a single H3 tail. Both mutants retain the ability to catalyze initial H3 mono-ubiquitination yet selectively decrease or enhance, respectively, the accumulation of multi-mono-ubiquitinated H3. The corresponding effects on DNA methylation maintenance therefore implicate the multi-mono-ubiquitinated state, rather than H3 mono-ubiquitination alone, in preserving methylation at replication-vulnerable CpGs. A likely mechanistic explanation is provided by the tandem UIMs within the DNMT1 RFTS domain, which engage multi-mono-ubiquitinated H3 through multivalent interactions and bind these substrates more strongly than singly ubiquitinated H3^32^. Increased avidity of DNMT1 for multi-mono-ubiquitinated H3 could therefore enhance its recruitment or retention at sites where maintenance methylation is most vulnerable to failure. This model is consistent with our previous finding that disruption of either DNMT1 UIM individually produces more modest methylation defects than disruption of both UIMs^35^, suggesting that the requirement for multivalent H3ub recognition becomes particularly important in genomic contexts where maintenance methylation is inefficient. Together, these observations provide complementary evidence from both sides of the UHRF1-H3ub-DNMT1 interaction that multi-mono-ubiquitination contributes functionally to maintenance methylation.

The selective requirement for UHRF1 ubiquitin reading at CpG-sparse regions further emphasizes that maintenance methylation mechanisms are strongly influenced by genomic context. Even in cells expressing WT UHRF1, a subset of CpGs progressively lost methylation during serial passaging, defining sites that are particularly vulnerable to replication-associated methylation erosion. Disruption of H3ub recognition accelerated methylation loss at these sites and expanded the population of vulnerable CpGs, whereas enhanced H3ub recognition by the DADA mutant partially buffered against this erosion. Importantly, these loci substantially overlap those previously shown to depend on UHRF1 E3 ligase activity^35^, providing convergent genetic evidence that both writing and subsequent recognition of H3ub contribute to maintenance methylation at CpG-sparse regions.

Why CpG-sparse regions are especially dependent on the UHRF1-H3ub-DNMT1 axis remains an important question. One possibility is that these regions provide fewer opportunities for DNMT1 to efficiently encounter hemi-methylated substrates during the limited post-replicative window for maintenance methylation, increasing the importance of mechanisms that retain or concentrate DNMT1 on newly replicated chromatin. Multi-mono-ubiquitinated H3 could provide such a mechanism by creating a multivalent chromatin-binding platform for the tandem UIMs of DNMT1. Consistent with this model, disruption of either UHRF1 ubiquitin writing or reading increased the RepliTali signature of replication-associated methylation loss. Progressive hypomethylation of late-replicating, CpG-sparse regions is a characteristic feature of both aging and cancer^11,12,35^, raising the possibility that reduced efficiency of the UHRF1-H3ub-DNMT1 pathway contributes to the gradual erosion of DNA methylation within these vulnerable genomic regions. Determining whether the activity or regulation of this pathway changes during aging or oncogenesis will be important for future testing of this hypothesis.

A notable limitation is that current approaches cannot comprehensively resolve the combinatorial ubiquitination state of individual H3 tails or map these states genome-wide. H3 multi-mono-ubiquitination has been identified by affinity-based enrichment and mass spectrometry^28,32^, but trypsin cleavage at H3R17 complicates determination of whether H3K14ub occurs on the same H3 molecule as H3K18ub or H3K23ub. Consequently, it remains unclear whether specific combinations of H3K14ub, H3K18ub, and H3K23ub are preferentially deposited at replication-vulnerable regions or confer distinct biological functions. Technologies capable of distinguishing single-from multi-mono-ubiquitinated H3 and mapping these combinatorial states to defined genomic loci will be important for determining how the spatial organization of H3 ubiquitination contributes to the fidelity of DNA methylation inheritance across different genomic contexts.

## Data availability

All EPIC array raw and processed data is deposited in the Gene Expression Omnibus archive under accession GSE347587 and will be released to the public upon publication.

## Supplementary data

Supplementary data are available at NAR online.

## Acknowledgements

We thank members of the Rothbart laboratory for helpful discussions. Computation for the work described in this paper was supported by the High-Performance Cluster and Cloud Computing (HPC3) Resource at the Van Andel Institute. We thank the Van Andel Institute Genomics Core (RRID:SCR_022913) for assistance with processing EPIC arrays. The graphical abstract was created in BioRender. Hrit, J. (2027) https://BioRender.com/d9h407w.

## Author contributions

Conceptualization: J.A.H., R.L.T., B.M.D., and S.B.R. Data curation: J.A.H. and R.L.T. Formal analysis: J.A.H., R.L.T., and B.M.D. Funding acquisition: J.A.H., R.L.T., and S.B.R. Investigation: J.A.H., R.L.T., V.J.S., and A.K.W. Methodology: J.A.H., R.L.T., and S.B.R. Project administration: S.B.R. Resources: J.A.H., R.L.T., A.K.W., E.J.W., and S.B.R. Software: J.A.H. and R.L.T. Supervision: E.J.W. and S.B.R. Validation: J.A.H., R.L.T., and S.B.R. Visualization: J.A.H. and R.L.T. Writing (original draft): J.A.H., R.L.T., V.J.S., and S.B.R. Writing (review and editing): all authors.

## Funding

National Institutes of Health [R35GM152184 to S.B.R., F32CA260116 to J.A.H., and R50CA315355 to R.L.T.]

## Conflict of Interest Statement

All authors declare no competing interests.

---

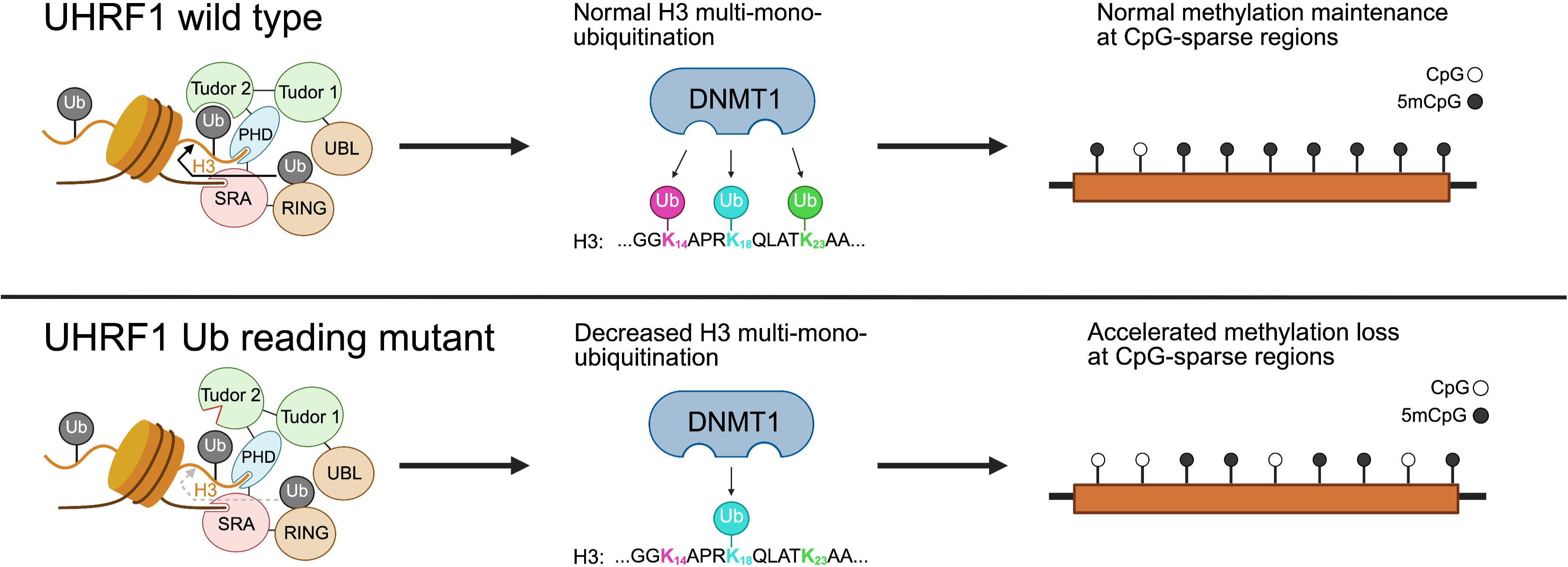

## Notes

### Competing Interest Statement

The authors have declared no competing interest.

