## Supplementary figures for "A feed-forward UHRF1 read-write mechanism supports H3 multi-mono-ubiquitination and DNA methylation maintenance at CpG-sparse regions"

### Figure S1

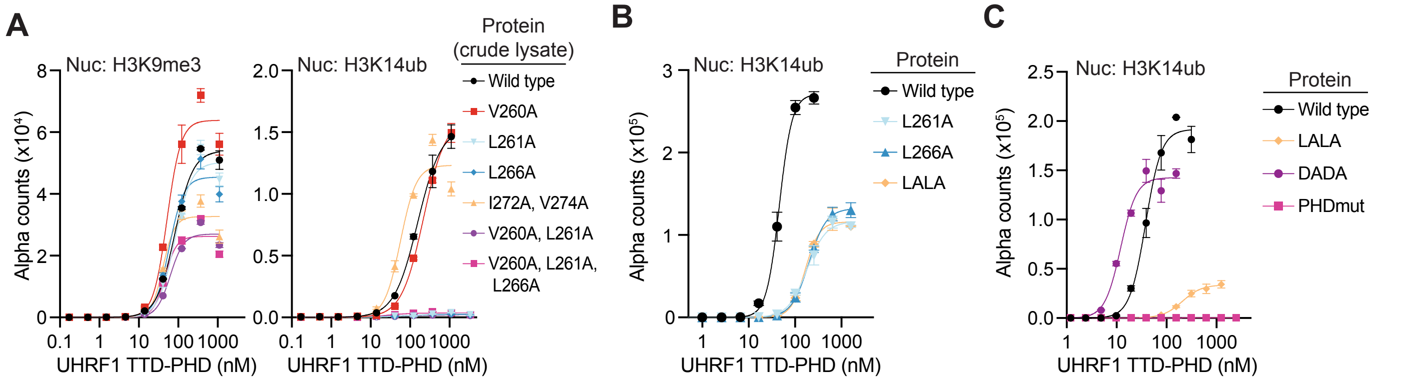


**D**

| UHRF1 TTD-PHD competitor | $IC_{50}$ (uM) vs. UHRF1 FL | $IC_{50}$ (uM) vs. RFTS |
|---|---|---|
| Wild type | 0.47 +/- 0.14 | 2.3 +/- 0.5 |
| LALA | n.m | n.m |
| DADA | 0.03 +/- 0.005 | 0.46 +/- 0.07 |
| PHDmut | n.m | n.m |

**Supplementary Figure 1 (related to Figure 2):**

(**A**) AlphaScreen binding assays measuring the interaction between *E. coli* lysates overexpressing wild-type or mutant UHRF1 TTD-PHD and the indicated biotinylated nucleosomes. Error bars represent SD of triplicate reactions. (**B-C**) AlphaScreen binding assays measuring the interaction between purified wild-type or mutant UHRF1 TTD-PHD and the indicated biotinylated nucleosomes. Error bars represent SD of triplicate reactions. (**D**) Table of $IC_{50}$ values measured in **Fig. 2F**. Error is +/- SEM.

Figure S2

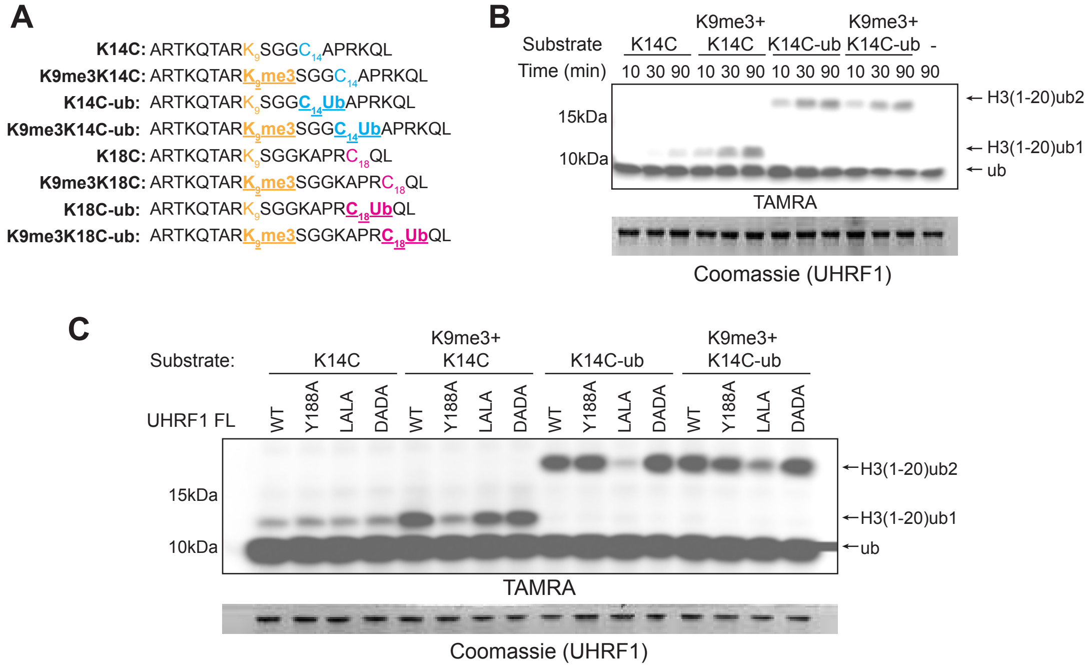
A
K14C: ARTKQTARK9SGGC14APRKQL
K9me3K14C: ARTKQTARK9me3SGGC14APRKQL
K14C-ub: ARTKQTARK9SGGC14UbAPRKQL
K9me3K14C-ub: ARTKQTARK9me3SGGC14UbAPRKQL
K18C: ARTKQTARK9SGGKAPRC18QL
K9me3K18C: ARTKQTARK9me3SGGKAPRC18QL
K18C-ub: ARTKQTARK9SGGKAPRC18UbQL
K9me3K18C-ub: ARTKQTARK9me3SGGKAPRC18UbQL
B
Substrate
K14C
K9me3+ K14C
K14C-ub
K9me3+ K14C-ub
-
Time (min)
10 30 90 10 30 90 10 30 90 10 30 90 90
15kDa
10kDa
H3(1-20)ub2
H3(1-20)ub1
ub
TAMRA
Coomassie (UHRF1)
C
Substrate:
K14C
K9me3+ K14C
K14C-ub
K9me3+ K14C-ub
UHRF1 FL
WT
Y188A
LALA
DADA
15kDa
10kDa
H3(1-20)ub2
H3(1-20)ub1
ub
TAMRA
Coomassie (UHRF1)

**Supplementary Figure 2 (related to Figure 3):**

(**A**) Semisynthetic H3 peptides used as substrates for UHRF1 ubiquitin ligase activity assays.
(**B-C**) *In vitro* ubiquitin ligase activity assays of recombinant UHRF1 using the indicated nucleosome substrates. TAMRA fluorescence shows ubiquitin and its products; Coomassie staining shows UHRF1.

### Figure S3

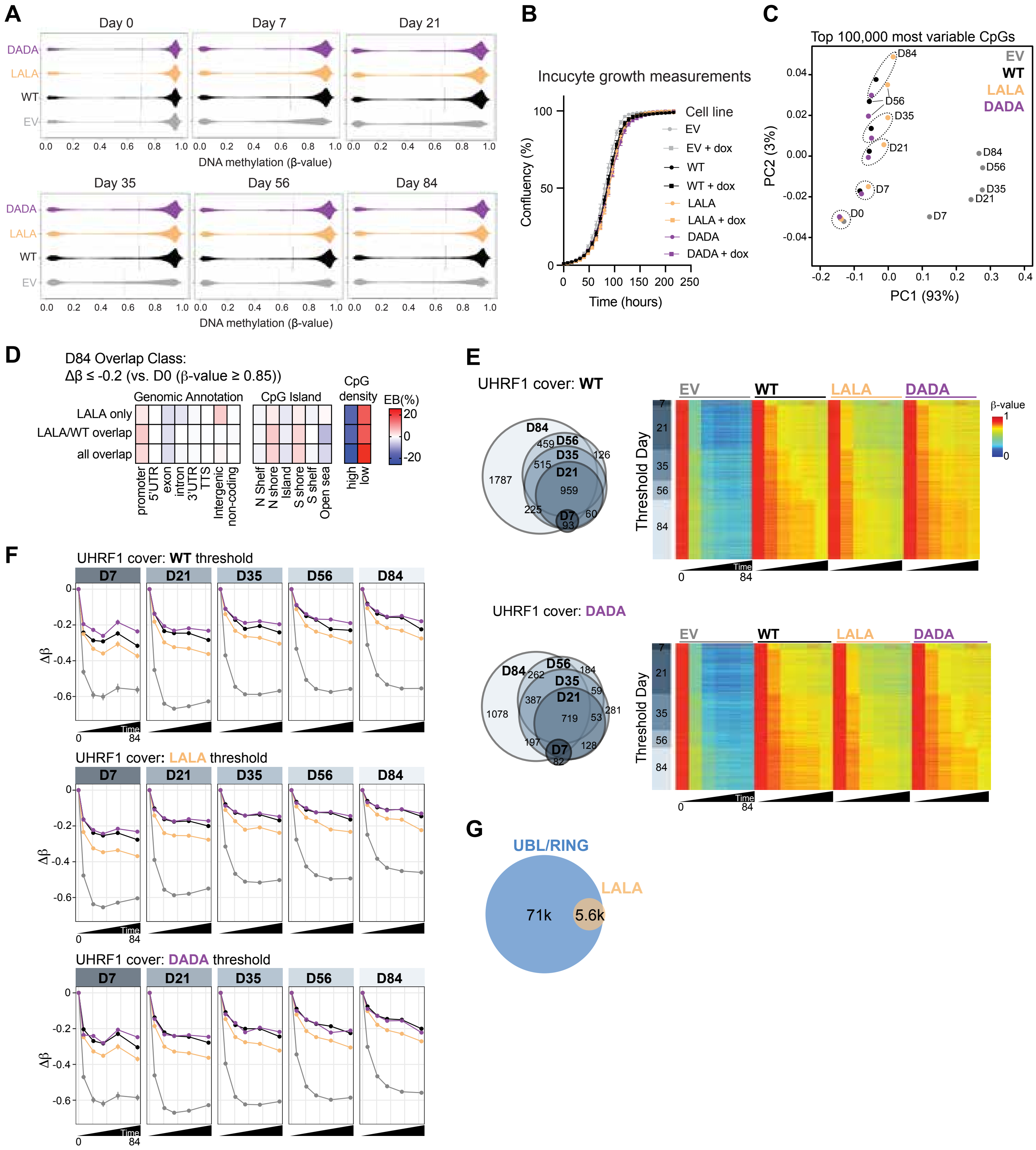

A
Day 0
Day 7
Day 21
DADA
LALA
WT
EV
DNA methylation (β-value)
Day 35
Day 56
Day 84
B
Incucyte growth measurements
Confluency (%)
Time (hours)
Cell line
EV
EV + dox
WT
WT + dox
LALA
LALA + dox
DADA
DADA + dox
C
Top 100,000 most variable CpGs
PC2 (3%)
PC1 (93%)
D
D84 Overlap Class:
Δβ ≤ -0.2 (vs. D0 (β-value ≥ 0.85))
Genomic Annotation
CpG Island
CpG density
EB(%)
LALA only
LALA/WT overlap
all overlap
promoter
5'UTR
exon
intron
3'UTR
TTS
Intergenic
non-coding
N Shelf
N shore
Island
S shore
S shelf
Open sea
high
low
E
UHRF1 cover: WT
UHRF1 cover: DADA
Threshold Day
β-value
Time
F
UHRF1 cover: WT threshold
UHRF1 cover: LALA threshold
UHRF1 cover: DADA threshold
Δβ
G
UBL/RING
LALA
71k
5.6k

**Supplementary Figure 3 (related to Figure 4):**

(**A**) Distribution of DNA methylation β-values across the indicated time points in HCT116 cells expressing EV, WT, LALA, or DADA transgene covers. (**B**) Incucyte growth measurements of cells expressing EV, WT, LALA, or DADA UHRF1 transgene covers in the presence or absence of doxycycline-induced depletion of endogenous UHRF1. (**C**) Principal component analysis of the 100,000 most variable CpGs across the experimental time course in cells expressing EV, WT, LALA, or DADA UHRF1 transgene covers. PC1 and PC2 explain 93% and 3% of the variance, respectively. (**D**) Enrichment of genomic annotations, CpG island context, and CpG density among probes classified by their day-84 hypomethylation overlap between LALA and WT UHRF1 cover conditions (related to **Figure 4E**). Probes were defined by $\Delta\beta \leq -0.2$ relative to highly methylated day 0 probes (β-value $\geq 0.85$) and classified as LALA-only, LALA/WT overlap, or all covers overlapping. Color indicates enrichment bias (EB%). (**E**) Temporal accumulation and DNA methylation dynamics of hypomethylated probes in cells expressing WT or DADA UHRF1 transgene covers. Venn diagrams show overlap among probes reaching the hypomethylation threshold at the indicated time points. Heatmaps show DNA methylation β-values across the 84-day time course in EV, WT, LALA, and DADA cells for probes grouped according to the first day at which the hypomethylation threshold was reached. (**F**) Average DNA methylation change (Δβ) across the 84-day time course for probes reaching the hypomethylation threshold at the indicated time points in WT-, LALA-, or DADA-expressing cells. Lines show methylation trajectories in EV, WT, LALA, and DADA conditions for each threshold-day probe group. (**G**) Venn diagram showing overlap of hypomethylated CpG probes ($\Delta\beta \leq -0.2$) that fail to maintain DNA methylation in cells expressing UHRF1 mutants affecting writer domains (UBL/RING) or H3ub reader activity (LALA).
